# Novel Behavioral Assays Reveal Sex Specific Behavioral Syndromes in Anemonefish

**DOI:** 10.1101/2024.06.07.597983

**Authors:** Gabriel J. Graham, Isabella M. Wilton, Emily K. Panczyk, Justin S. Rhodes

## Abstract

Sexually dimorphic behaviors are common across taxa, particularly in the contexts of parental care and territorial aggression. The false clown anemonefish *Amphiprion ocellaris* is unique among animals for its combination of female behavioral dominance and territoriality, protandrous sex change, and mutualistic symbiosis with sea anemones. Several laboratory studies have begun characterizing sex differences in parental care and aggression in this species, but aggression assays have mostly focused on intra-specific aggression where individual differences are large. The goals of this study were to expand the repertoire of behavioral assays available for *A. ocellaris*, establish repeatability of individual differences, identify assays that produce the most robust sex differences, and explore whether individual differences in correlated behaviors can be detected consistently across experimental contexts (i.e., whether behavioral syndromes can be detected). To this end, we measured 39 behaviors across 7 behavioral assays (parental care, large intruder aggression, small intruder aggression, male-oriented aggression, female-oriented aggression, immediate reaction to a threat, and nest maintenance) in 9 reproductively active *A. ocellaris* pairs under 3 different contexts (without eggs in the nest, with their own eggs, and with surrogate eggs). Behaviors were repeatedly measured three separate times (rounds) over repeated spawning cycles. We found 34 out of 39 behaviors were significantly individually repeatable across egg contexts and rounds, with an average intra-class correlation of 0.33. We found parental care, large intruder aggression, and female-oriented aggression assays produced the largest sex differences. Males performed 7-fold more egg care behaviors than females, while females performed 3.5-fold more aggressive behaviors toward a large interspecific intruder. Further, females bit, chased and struck the female intruder 6.5, 4.1, and 3.2 times as many times as males did. Five different behavioral syndromes were observed in males but only one was observed in females. These results expand our understanding of sex differences in behavior and the division of labor in the iconic anemonefish. Future studies can use these assays to measure the behavioral sex of fish in the middle of sex change, in the study of behavioral plasticity, or in the study of the neuroendocrine bases of aggression and parental care in this unique species.

## Introduction

Behavioral sexual dimorphisms are common in sexually reproducing species. Species that care for their young and defend a territory often display a division of labor. For example, in many species (e.g., the African cichlid *Neolamprologous pulcher*, the marine goby *Microgobius gulosus*), males defend the territory while females care for eggs. The false clown anemonefish *Amphiprion ocellaris*, however, defies this norm. In this iconic coral reef fish, males are the primary caregivers, while females are responsible for territory defense. While paternal care is not uncommon in fish (e.g., in damselfishes, *Dascyllus reticulatus* and *Dascyllus aruanus,* and gobies *Pomatoschistus minutus* and *Paragobiodon echinocephalus*) (Gross & Sargent, 1985; Kuwamura et al., 1993; Lindström et al., 2006; Mizushima et al., 2000; Sakai et al., 2024), it is relatively unusual for females to take the primary role in territory defense (Kölliker, 2012). Even more unusual, anemonefish undergo protandrous (male-to-female) sex change (Fricke & Fricke, 1977; Liu et al., 2017). In the wild, *A. ocellaris* live in small, isolated groups comprised of one reproductively active α female, one reproductively active β male, and potentially several non-reproductive γ males (Fricke & Fricke, 1977). Should the α female be killed or removed, the β male takes her spot in the dominance hierarchy and undergoes protandrous sex change to become the new α female, while the most dominant γ male ascends to become the β male, and all subsequent fish move up one position in the dominance hierarchy (Fricke & Fricke, 1977). Thus, the same individual can display both male and female behaviors during their lifetime.

Anemonefish are also unique in that they are entirely dependent on their host sea anemone for survival (Da Silva & Nedosyko, 2016). Because of this dependence, they display extremely high levels of territorial defense (Laudet & Ravasi, 2022). Anemonefish will stand their ground and fight when faced with a significantly larger threat while most other coral reef fish species will swim away and hide in the rocks of the reef. As the largest and behaviorally dominant member of the group, females are the most aggressive (Iwata & Manbo, 2013). For example, female anemonefish have been known to attack much larger foes including human divers and sharks that stray too close to their host sea anemone (Iwata & Manbo, 2013; Laudet & Ravasi, 2022). This makes anemonefish ideal for studying female behavioral dominance and high levels of territorial aggression in females (Iwata & Manbo, 2013; Laudet & Ravasi, 2022; Rhoades & Blumstein, 2007).

Anemonefish’s dependence on their sea anemone also means that they are particularly well suited to laboratory research. In the wild, anemonefish rarely stray more than a few meters away from the safety of their sea anemone, thus, they are comfortable in the confines of small (∼75L) aquaria (Laudet & Ravasi, 2022). Additionally, in the lab, anemonefish readily reproduce and display normal parental care and nest behaviors even in the absence of a sea anemone (R. DeAngelis et al., 2017, 2020; Laudet & Ravasi, 2022; Phillips et al., 2020).

Recent laboratory studies have uncovered an unusual feature of anemonefish parental care behavior that allows for useful manipulations that are not possible for most other species. Anemonefish will care for eggs that are not their own and not kin with similar vigor as they would their own eggs (Phillips et al., 2020). Thus, parental care behavior can be evaluated by presenting fish with eggs from a different spawning pair and recording various parental care behaviors. This is a relatively unusual feature in fish as most fish will eat eggs that are not closely related (Pereira et al., 2017). The ability to use surrogate eggs can be useful for evaluating parental care behaviors in fish that may not have their own eggs (such as fish in the middle of sex change or in newly formed pairs) (Parker et al., 2022).

Recent laboratory studies have also begun to evaluate differences in levels of parental care and aggression displayed by male and female anemonefish (R. DeAngelis et al., 2020; Iwata & Manbo, 2013; Phillips et al., 2020). While males and females on average display different levels of parental care and territorial aggression, there is substantial individual variation that can obscure sex differences (R. S. DeAngelis & Rhodes, 2016; Iwata & Manbo, 2013; Parker et al., 2022; Phillips et al., 2020). Males also defend the nest and females also care for eggs, just to a lesser extent. The laboratory assays that have been used to establish differences in aggression in *A. ocellaris*, have mostly focused on intra-specific aggression toward an intruder male or female (Iwata & Manbo, 2013; Parker et al., 2022; Yaeger et al., 2014). In the assays used in Parker et al. (2020), males and females were removed from their home tank and paired with a novel male or female in a neutral territory. A sex difference was discerned using this assay, especially when considering the ratio of female-oriented to male-oriented aggression. Females tend to be more aggressive to females and less aggressive to males whereas males show the reverse pattern. However, there is substantial overlap in the distributions even for the ratio (Parker et al., 2022). Thus, additional work is needed to broaden the repertoire of behaviors to find the most robust, consistent assays that are demonstrative of the differential roles each sex has in the reproductive pair, especially for territorial aggression toward non-conspecifics. This is important not only for documenting the life history of an iconic species of fish but also useful for mechanistic studies attempting to uncover neurological or neuroendocrine regulation of sex differences in parental care and territorial aggression (R. DeAngelis et al., 2018, 2020; R. S. DeAngelis & Rhodes, 2016; Iwata & Manbo, 2013; Laudet & Ravasi, 2022).

Additionally, while maintaining aquaria, we have noticed pronounced personalities among various anemonefish. When approaching an aquarium for basic maintenance, some females are passive and hide in the back of the tank, while others are extremely aggressive, and leap out of the water in an attempt to bite us. Moreover, some males display substantially more effort in caring for eggs than others (unpublished observations). However, to our knowledge, no one has evaluated the extent to which repeatable individual differences in aggressive and parental behaviors can be detected in the lab, i.e., whether laboratory housed anemonefish display behavioral syndromes, personalities, or reliable correlated sets of behaviors.

The goal of this study was twofold: First to find behavioral assays that most reliably differentiate the sexes and second to identify repeatable individual differences and examine if these individual differences constitute behavioral syndromes: correlated sets of behaviors that are repeatable across various environmental contexts (Bell et al., 2009).

## Methods

### Animals and Husbandry

Anemonefish used in this study were either first or second generation bred in-house from fish originally obtained from Ocean Reefs and Aquariums (Fort Pierce, FL), obtained as a gift from Peter Buston, or purchased from the pet trade. Fish were housed in 20-gallon tall (24” x 12” x 16”) or 25-gallon cube (18” ×18” ×18”) centrally filtered aquariums with a 6” terra cotta pot serving as a nest site and spawning substrate.

The Domino damselfish (*Dascyllus trimaculatus*) and hermit crabs (*Coenbita spp.*) used in this study were obtained from the pet trade and were housed together in 10-gallon (20” x 10” x 12”) aquariums consisting of three damselfish and one hermit crab. Filtration was provided by sponge filters with reef rocks and terra cotta pots serving as cover.

In all tanks, salinity was kept at 1.026 specific gravity, temperature was kept between 26°-28°C, pH was kept between 8.0 and 8.4, and lighting was provided by fluorescent and LED lights over tanks on a 12:12 light schedule (lights on at 07:00 h and off at 19:00 h). Fish were fed twice daily with ReefNutrition TDO Chroma Boost pellets (Reed Mariculture Inc., Campbell, CA).

### Ethical Note

None of the species used in this study are protected or endangered. All experimental procedures were approved by the University of Illinois Institutional Animal Care and Use Committee.

### Behavioral Assays

We performed 7 behavioral assays to study the behaviors of 9 reproductively active pairs (n=18, 9 males and 9 females; see Supplementary Excel File 1 for weights and lengths of animals). All assays were recorded between 15:00 and 17:00 to control for possible effects of the circadian rhythm on the behaviors. For the purposes of this study, the term **focal animal** signifies the fish whose behavior was recorded, and the term **stimulus animal** signifies the animal that was introduced into the tank. After recording began, the researcher left the area to avoid influencing the behavior of the animals. We chose to use the same assay order across all conditions to keep possible order effects consistent in our analyses. Across all egg conditions and rounds, the behavioral assays were performed in the following order:

1. <u>Parental Care.</u> We used a variant of the parental care assay from in Phillips et al. (2020), Parker et al. (2022), R. DeAngelis et al., (2020), DeAngelis et al. (2018), and DeAngelis & Rhodes (2016). Males and females were recorded for 5 minutes with eggs in the nest, and for each individual, the following parameters were measured: nips, pectoral fans, tail fans, nest entrances, and time in the nest (See table 1 for descriptions of behaviors). In the no eggs condition, this assay was omitted.
2. <u>Small Intruder Aggression.</u> The day following the parental care assay, we tested the focal animals’ reaction to a small intruder by placing a hermit crab near the nest. A total of 4 hermit crabs were used in this assay and rotated randomly across the various trials. To control movement around the tank, we fashioned a “leash” out of a centrifuge tube glued to the crab’s shell with a small stretch of fishing line connecting it to another centrifuge tube that was placed under the terra cotta pot. This prevented the crab from moving more than approximately a 10 cm radius away from the nest. Focal animals’ behaviors were recorded for 10 minutes. We measured the latency to the first recorded behavior, the number of bites, and strikes over the 10-minute period.
3. <u>Large Intruder Aggression.</u> The day after the small intruder assay, we tested the focal animals’ reaction to a large intruder in the tank. Domino damselfish (*Dascyllus trimaculatus*) were selected since they are commonly found in close proximity with *A. ocellaris* in nature and are considered competitors and potential egg predators (Hayashi et al., 2020). This assay was modified from the assay used in DeAngelis et al. (2020). Here, we only used one damselfish as opposed to the three used in the cited study to allow for more targeted aggression. The stimulus animal was placed into the tank for 10 minutes and the following behaviors were recorded: latency to first recorded behavior, and the number of bites, strikes, displays, and chases/charges. A total of 9 different individual damselfish were used in this assay and rotated randomly across the different trials. Damselfish were 3.5 to 4.5 cm in length.
4. <u>Male-Oriented Aggression.</u> The day following the large intruder assay, we tested the focal animals’ reaction to a novel conspecific male. This assay along with the female-oriented aggression assay were modified from the assays used in Parker et al. (2022) in two relevant ways. First, in this study, the focal pair was kept together and second, the assay took place in their home tank as opposed to a novel tank. Due to high levels of aggression from some fish, the male was only placed in the tank for 2 minutes as any longer was deemed to be a risk to the stimulus animal’s health. The following behaviors were recorded: latency to first recorded behavior, and number of bites, strikes, displays, and chases/charges. A total of 29 different stimulus males were used in this assay and rotated randomly across the different trials (see Supplementary Excel File 1 for weights and lengths of males).
5. <u>Female-Oriented Aggression.</u> The day after the male-oriented aggression assay, we tested the focal animals’ reaction to a conspecific female. Only females that had previously spawned were used, and we recorded the stimulus females’ relative size to the focal female to use as a covariate. For similar reasons as the male-oriented aggression assay, the stimulus animal was only placed in the tank for 2 minutes. The following behaviors were recorded: latency to the first recorded behavior, bites, strikes, displays, chases/charges. A total of 28 different stimulus females were used in this assay and rotated randomly across the different trials (see Supplementary Excel File 1 for weights and lengths of females).
6. <u>Immediate Reaction to a Threat.</u> The day after the female-oriented aggression assay, we tested the focal animals’ immediate reaction to a large threat, a proxy for boldness. To test this, a researcher placed their hand in the aquarium and behaviors were recorded for 10 seconds. The following behaviors were recorded: latency to first recorded behavior, and number of bites, strikes, displays, and chases/charges.
7. <u>Nest Maintenance.</u> The same day as the immediate reaction to a threat assay (at least 10 minutes after), we tested how the focal pair would respond to a nest contaminant. We placed a small lava rock (∼ 4 cm.) in the back of the terra cotta pot and recorded the following behaviors for 5 minutes: latency to the first recorded behavior, and number of bites, and strikes. On a few occasions, the pair would successfully remove the rock from the pot prior to the end of the 5 minutes. In this case, the assay was redone the following day.

### Egg Conditions

To determine how the various behaviors measured in the behavioral assays were impacted by the presence of eggs in the nest, we repeated all the behavioral assays across three egg conditions. Assays were recorded when the sample pair had their own eggs (Own), surrogate eggs of a similar age from a different pair (Surrogate), and no eggs (None / No). In the no eggs condition, the parental care assay was omitted. To control for hormonal changes across the breeding cycle, surrogate eggs were exchanged with the pair’s own eggs the day after spawning such that the pair would be in the same position in its breeding cycle in both conditions (R. S. DeAngelis & Rhodes, 2016). When exchanging surrogate eggs for the pair’s own eggs, 24h was allowed for acclimation to the novel terra cotta pot (Unpublished observation, anemonefish require 24h of acclimation to behave normally in the surrogate condition and for behaviors to stabilize). The no egg condition began the day after all eggs had naturally hatched or died, and if the pair spawned again before all assays could be completed, the assays were completed in the next inter-spawn period.

### Rounds

To measure the repeatability of the behavioral assays, we repeated all the assays in each egg condition three times during three separate spawning cycles.

### Behavioral Analysis

All assays were recorded using Samsung HMX-W300 cameras. 24 Hours prior to the first assay until the end of the last behavioral assay, the camera and tripod were set up in front of the tank to acclimatize the pairs to recording. Videos were analyzed using BORIS (v. 8.22) to record and timestamp the behaviors as the video progressed (Friard & Gamba, 2016). The timer for each analysis began after the researcher’s hand left the tank or the stimulus was added and ended after the allotted amount of time had passed. Behaviors were only recorded in this time window.

#### Parental Behaviors

**Table 1A.** Descriptions of parental behaviors.

|  |  |
| --- | --- |
| Time in nest | More than half of the focal animal's body is inside the terra cotta pot. |
| Entrance | The fish swims into the terra cotta pot. |
| Pectoral fan | The focal animal waves its pectoral fins at the eggs. |
| Tail fan | The focal animal 'wiggles' its tail fin at the eggs. |
| Nip | The focal animal mouths/ bites at the eggs. |

#### Aggressive Behaviors

**Table 1B.** Descriptions of aggressive behaviors.

|  |  |
| --- | --- |
| Latency | The time from the start of the video to the first recorded behavior. If no behavior was observed, the length of the assay was counted as the latency. |
| Chase/Charge | The focal animal swims quickly at the stimulus animal. |
| Display | The focal animal shows its side to the stimulus animal. This may include floating at an angle, tilting its body, or flexing its body. |
| Strike | The focal animal bats the stimulus animal with its tail or rams the stimulus animal with its head. |
| Bite | The focal animal bites the stimulus animal. |

The terms **sum of parental behaviors, sum of aggressive behaviors,** or **sum of behaviors** indicate the sum of measured behaviors in each assay not including latency. In the parental care assay, nest entrances and time in nest do not contribute toward this sum.

#### Aggression Ratio

As in Parker et al. (2022), a female to male aggression ratio was calculated. The sum of aggressive behaviors in the female-oriented aggression assay was divided by the sum of aggressive behaviors in the male-oriented aggression. We added a value of 1 to both of these values to avoid zeros in the denominator. The female/male aggression ratio was then log10 transformed due to non-normal distribution of residuals. This results in a measure centered around 0 where positive values indicate a female aggression bias, and negative values indicate a male aggression bias.

### Statistical Analyses

R (4.2.3) was used for statistical analyses with a *P*< 0.05 considered statistically significant (R Core Team, 2024).

#### Linear Regression

The lmer() function from the package “lme4” was used to fit linear mixed-effects models to the behaviors in each assay. We modeled both the sum of behaviors recorded in the assay and the count of each individual behavior within each assay. Fixed effects included round, egg condition and sex with sex by round and sex by egg condition interactions. Note that round and egg condition are within-subjects fixed effects whereas sex is between-subjects. The fish id was entered as a random effect to account for repeated measures. In several analyses we included covariates as predictor variables in the models. In the female-oriented aggression assay, we included the difference in size between the focal female and the stimulus female. We evaluated a similar covariate for the male-oriented assay, the difference in size between the focal male and the stimulus male, but since it was not significant in any of the analyses it was omitted from the final analyses. In analyses where we compared own eggs to surrogate eggs groups, we included the age of the eggs and the number of eggs in the nest as covariates. In instances where residuals were not normally distributed, we transformed data with an appropriate transformation (see Supplementary Excel Files). In cases where we log transformed data, we added 1 to all values to avoid taking the log of zero which is undefined. Whenever we refer to fold-changes throughout the manuscript, the calculation is done on raw untransformed values. To extract P-values for individual factors with multiple levels (e.g., round, egg condition), we used the Anova() function from the package “car” with the type = “III” option specified. Pairwise comparisons were computed using the emmeans() functions from the package “emmeans” with the tukey P-value adjustment specified. Partial eta squared (η^2^_p_) for mixed effects models and eta squared (η^2^) for linear regression models were calculated using the eta_squared() function from the package “effectsize”.

#### Repeatability

To estimate repeatability of each behavioral measure, we calculated the intra-class correlation (ICC). To do this, we divided the estimate of the individual variance from the mixed effects linear model (that included round, egg condition, sex, sex by round and sex by egg condition interactions as factors; see above) by the sum of residual variance + individual variance. To calculate the significance of repeatability, we extracted the significance of the random effect using the function ranova() from the package ‘lmerTest’.

#### Principal Component Analysis

The princomp() function was used to perform principal component analyses. Two types of analyses were performed. The first analyzed the behavior of each fish across all 7 assays by using the sum of behaviors collapsed across the egg conditions and rounds for each behavioral assay as input variables. The second analyzed the behavioral composition within each assay by using each of the recorded behaviors as input variables. The cor = TRUE option was specified to correct for differences in the scale of the measurements within and between assays. For example, in the analysis that included all behavioral assays, some of the assays were sums of behaviors collected over 10 seconds (e.g., immediate reaction to a threat) whereas others occurred over 5 minutes (e.g., nest maintenance). Sex differences in PC1 and PC2 were evaluated using unpaired t-tests.

## Results

### Parental Care: Males contribute significantly more time and effort toward parental care than females

We began our experiment by using an assay that is well known to produce large and significant differences between males and females: a parental care assay (R. DeAngelis et al., 2017, 2018; R. S. DeAngelis & Rhodes, 2016; Parker et al., 2022; Phillips et al., 2020). As expected, the sum of parental behaviors was highly sexually dimorphic (Square root transformed, χ^2^_1_= 18.060, *P*< 0.001, η^2^_p_= 0.725; Fig. 1A), as well as time spent in the nest (No transformation, χ^2^_1_= 9.770, *P*< 0.001, η^2^_p_= 0.619; Fig. 1B). Males spent over 3 times as much time in the nest and produced over 7 times as many parental care behaviors compared to females. Considering each behavior individually, nips (Square root transformed, χ^2^_1_= 8.319, *P*= 0.004, η^2^_p_= 0.588; supplementary Fig. 1A), and pectoral fans (Square root transformed, χ^2^_1_= 19.943, *P*< 0.001, η^2^_p_= 0.737; supplementary Fig. 1B), were significantly different between males and females, but not tail fans (*P*>0.05), likely due to their rarity. As in Phillips et al. (2020), we found that pairs behaved similarly regardless of whether they were caring for their own eggs or surrogate eggs (All *P*>0.05; see Supplementary Excel File 2 for ANOVA tables, and 3 for means and standard errors).

**Figure 1.**
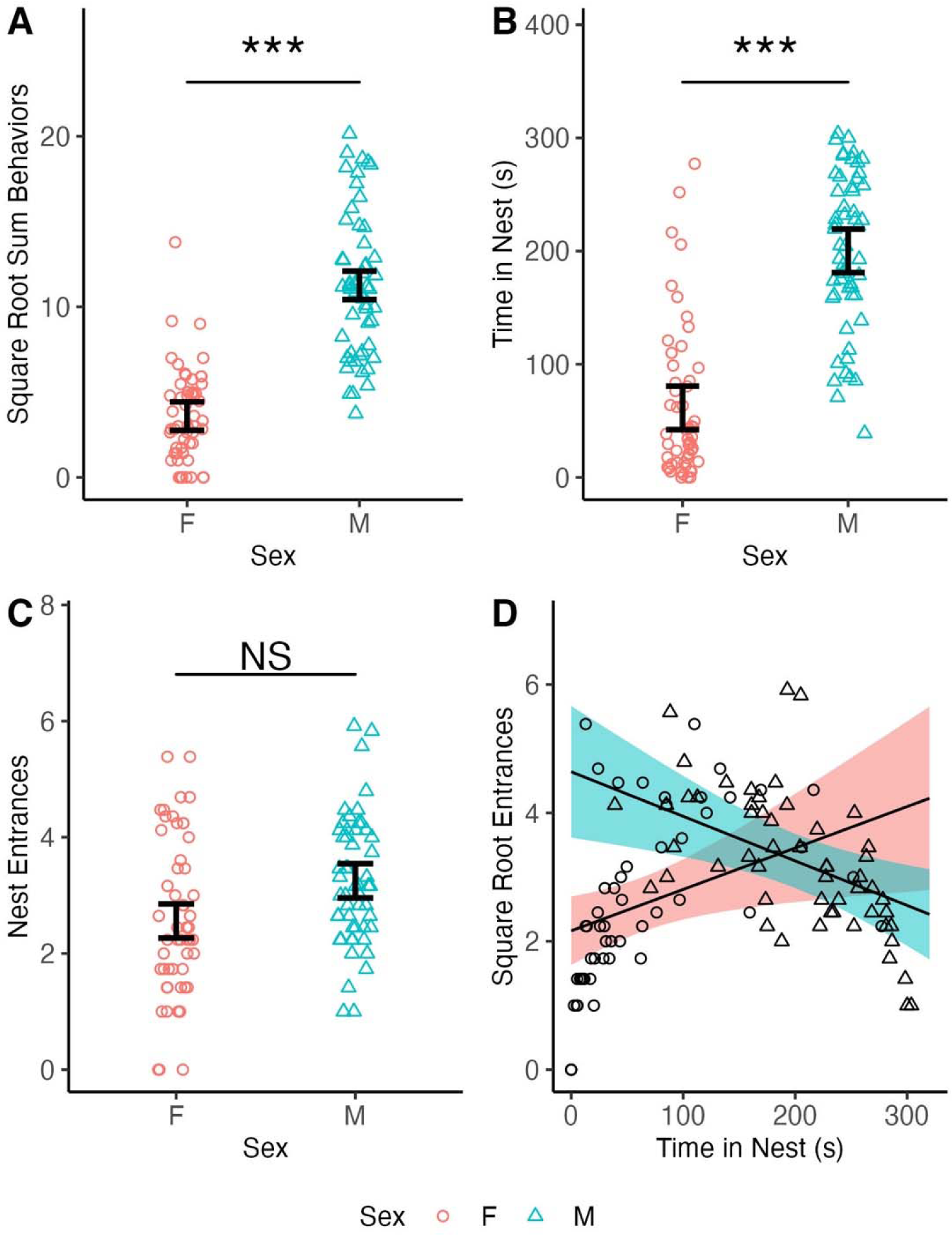
Males are the primary caregivers of the eggs. All data points are shown including repeated measures across the 2 egg conditions and 3 rounds. A) The square root of the sum of parental behaviors by sex. Males performed 7-fold more parental behaviors than females. B) The amount of time spent in the nest by sex. Males spent 3-fold more time in the nest than females. C) The square root of the number of nest entrances by sex. Males entered the nest slightly more than females, but not significantly. D) The square root of entrances plotted against time spent in nest, split by sex. There was a significant square root entrances by time interaction (*P*< 0.001); males show a negative correlation whereas females show a positive correlation. Lines represent linear regression estimates and ribbons represent the ± 95% confidence interval from the mixed model. In A-C, error bars show the standard error estimate from the linear mixed model that represents individual variance centered around the mean estimate from the model, *** indicates *P*< 0.001, and “NS” indicates not significant.

In addition to parental care behaviors and time in the nest, we also measured the number of nest entrances. While the sexes did not differ in the number of nest entrances (*P*> 0.05, Fig. 1C), the relationship between time spent in the nest and number of nest entrances was opposite for males as compared to females (i.e., the interaction between sex and time in the nest for square root transformed number of entrances was significant; Square root transformed, χ^2^_2_= 14.564, *P*< 0.001, η^2^_p_= 0.200; Fig. 1D). Males spent less time in the nest as nest entrances increased, while females spent more time in the nest as nest entrances increased. This reverse pattern is likely a reflection of the division of labor regarding parental care and territorial defense. Males spend most of their time caring for eggs and only rarely leave the nest to defend the territory or explore the environment, while females spend the majority of their time out of the nest exploring the environment. Thus, in males, an increase in nest entrances indicates increased time out of the nest relative to other males, while in females, an increase in nest entrances indicates increased time in the nest as compared to other females (see Supplementary Excel File 2 for ANOVA tables, 3 for means and standard errors, and 4 for pairwise comparisons).

All recorded parental care behaviors were significantly repeatable (see table 2 for ICC and P-values).

**Table 2.**
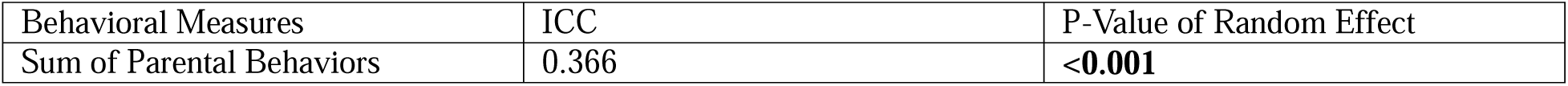
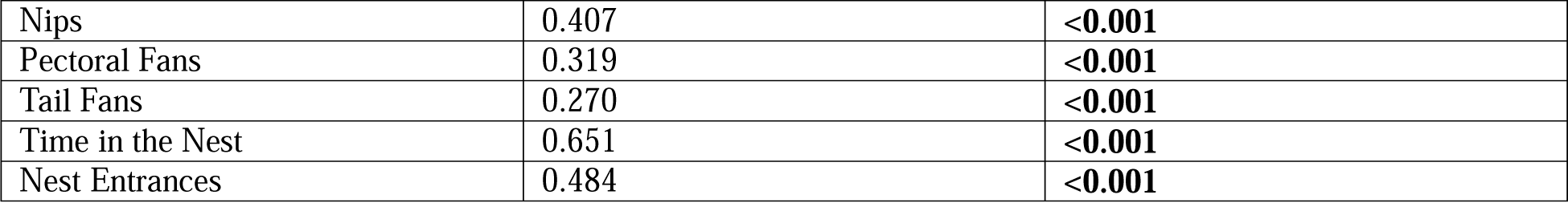
ICC and P-value for the random effect for all measured behaviors in the parental care assay.

| Behavioral Measures | ICC | P-Value of Random Effect |
| --- | --- | --- |
| Sum of Parental Behaviors | 0.366 | **<0.001** |
| Nips | 0.407 | **<0.001** |
| Pectoral Fans | 0.319 | **<0.001** |
| Tail Fans | 0.270 | **<0.001** |
| Time in the Nest | 0.651 | **<0.001** |
| Nest Entrances | 0.484 | **<0.001** |

### Small Intruder Aggression: Pairs are more aggressive when they have eggs in the nest

A hermit crab was placed near the nest to measure the pairs’ response to a small intruder. Females produced 1.5-fold more strikes than males (Square root transformed, χ^2^_1_= 4.879, *P*= 0.027, η^2^_p_= 0.149; Fig. 2A; see Supplementary Excel file 5 for ANOVA tables and 6 for means and standard errors). No significant effects of egg condition, round, or any interactions were detected. None of the other measured behaviors in this assay showed significant effects of sex, egg condition, round or interactions (All *P*>0.05).

**Figure 2.**
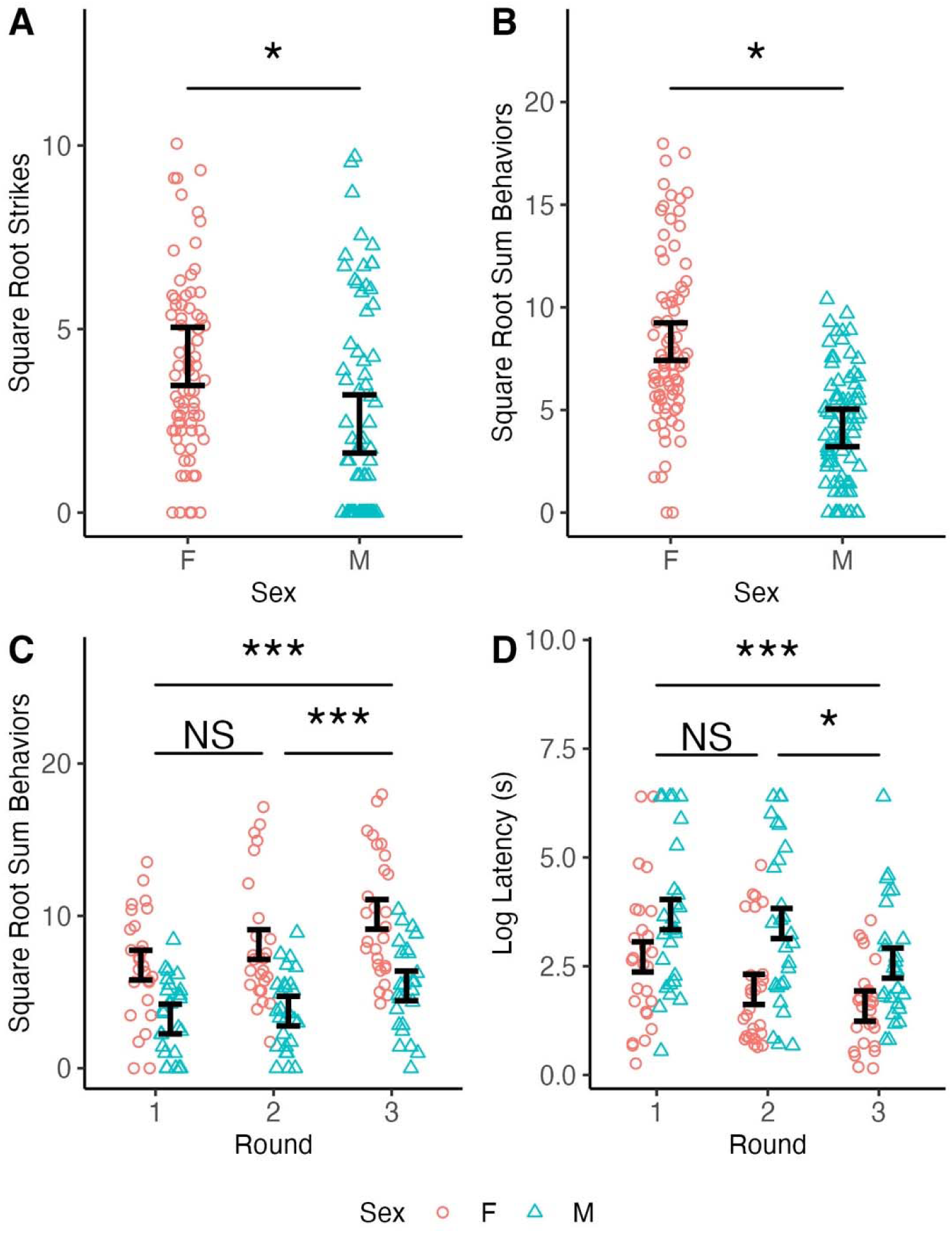
Females are more aggressive toward intruders than males, and aggression toward a large intruder increases as rounds progress. All data points are shown including repeated measures. A) The square root of strikes in small intruder aggression shown by sex. Females struck the small intruder 1.5 times more than males. B) The **s**quare root of sum of aggressive behaviors toward the stimulus animal in large intruder aggression shown by sex. Females displayed 3.5-fold more aggressive behaviors than males. C) The square root of the sum of aggressive behaviors in large intruder aggression shown by round and sex. Pairs got more aggressive as rounds progressed. D) The log of latency to address the stimulus animal in large intruder aggression by round and sex. Pairs addressed the intruder more quickly as rounds progressed. Error bars show the standard error estimate from the linear mixed model that represents individual variance centered around the mean estimate from the model, *** indicates *P*< 0.001, * indicates *P*< 0.05, and “NS” indicates not significant.

All recorded small intruder behaviors were significantly repeatable (see table 3 for ICC and P-values).

**Table 3.** ICC and P-value for the random effect for all measured behaviors in the small intruder aggression assay.

| Behavioral Measure | ICC | P-Value of Random Effect |
| --- | --- | --- |
| Sum of Aggressive Behaviors | 0.664 | **<0.001** |
| Latency | 0.458 | **<0.001** |
| Strikes | 0.662 | **<0.001** |
| Bites | 0.462 | **<0.001** |

### Large Intruder Aggression: Females contribute more toward territory defense

To model the fish’s response to a large intruder, we introduced a domino damselfish *(Dascyllus trimaculatus*) into the home tank of the focal pair and measured their behavioral responses.

We found a significant effect of sex on the sum of aggressive behaviors (Square root transformed, χ^2^_1_= 4.968, *P*= 0.026, η^2^_p_= 0.397; Fig. 2B); females produced more than 3.5 times the number of aggressive behaviors than males (see Supplementary Excel File 7 for ANOVA tables, and 8 for means and standard errors). No significant effects of egg condition were detected for the sum of aggressive behaviors or for any of the behaviors individually (All *P* >0.05).

We did however find a significant effect of round on the sum of aggressive behaviors (Square root transformed, χ^2^_2_= 33.070, *P*< 0.001, η^2^_p_= 0.253; Fig. 2C), chases and charges (Log transformed, χ^2^_2_= 12.477, *P*= 0.002, η^2^_p_= 0.135; supplementary Fig. 1C), displays, (Square root transformation, χ^2^_2_= 24.236, *P*< 0.001, η^2^_p_= 0.200; supplementary Fig. 1D), and latency (Log transformed, χ^2^_2_= 9.879, *P*= 0.007, η^2^_p_= 0.122; Fig. 2D). In all of these cases, pairs increased in aggression as rounds progressed as indicated by a 2-fold increase in the number of aggressive behaviors including a 2.5-fold increase in chases/charges, a 1.8-fold increase in displays, and a 77% reduction in latency between round 1 and round 3 (see Supplementary Excel File 4 for pairwise comparisons, 7 for ANOVA tables, and 8 for means and standard errors).

All recorded large intruder behaviors were significantly repeatable (see table 4 for ICC and P-values).

**Table 4.** ICC and P-values for the random effect for all measured behaviors in the large intruder aggression assay.

| Behavioral Measure | ICC | P-Value of Random Effect |
| --- | --- | --- |
| Sum of Aggressive Behaviors | 0.605 | **<0.001** |
| Latency | 0.211 | **<0.001** |
| Displays | 0.808 | **<0.001** |
| Chases/Charges | 0.240 | **<0.001** |
| Strikes | 0.181 | **<0.001** |
| Bites | 0.136 | **0.008** |

### Male-Oriented Aggression: Pairs get more aggressive toward male intruders over time

For the male-oriented aggression assay, we introduced a novel stimulus male into the home tank of the focal pair and measured their behavioral responses. Both focal males and females displayed similar levels of aggression toward the stimulus male (All *P* >0.05).

However, we did find a significant effect of egg condition on latency (Log transformed, χ^2^_2_= 16.626, *P*<0.001, η^2^_p_= 0.095; Fig. 3A). Interestingly, pairs addressed the stimulus male approximately 52% faster in the no egg or surrogate eggs condition compared to the own eggs condition, a significant difference, but did not significantly differ when comparing no eggs to surrogate eggs (see Supplementary Excel File 4 for pairwise comparisons, 9 for ANOVA tables, and 10 for means and standard errors).

**Figure 3.**
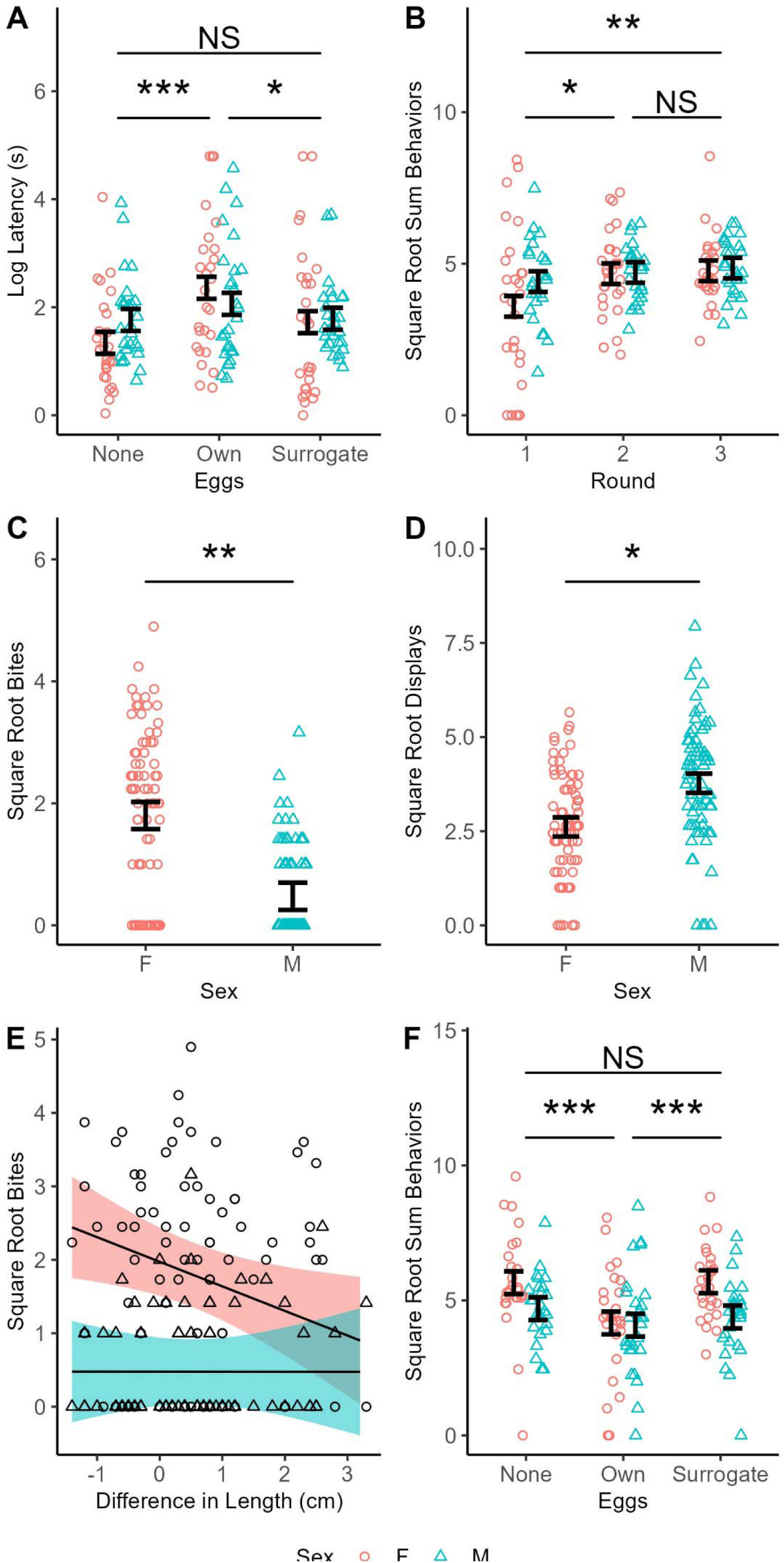
Females are more aggressive toward female intruders than males, but both sexes are similarly aggressive toward male intruders. All datapoints are shown including repeated measures. A) The log of latency to address the stimulus male in male-oriented aggression shown by egg condition and sex. Pairs responded to the stimulus male 50% more slowly when they had their own eggs in the nest as compared to when they had surrogate eggs or no eggs. No sex differences were detected. B) The square root of the sum of aggressive behaviors toward the stimulus male in male-oriented aggression assay shown by round and sex. Pairs displayed 10% more aggressive acts in rounds 2 and 3 as compared to round 1. No sex differences were detected. C) The square root of bites toward the stimulus female in female-oriented aggression shown by sex. Females bit the female stimulus animal 6.5-fold more times than males. D) The **s**quare root of displays toward the stimulus female in female-oriented aggression shown by sex. Males displayed toward the stimulus female 1.9-fold more than females. E) The square root of bites in female-oriented aggression shown by the difference in length between the focal female and stimulus female, and sex. For each centimeter increase in the difference in length between the focal female and stimulus female, pairs get significantly (*P*= 0.008) less aggressive, particularly females. Lines represent linear regression estimates and ribbons represent the ± 95% confidence interval from the mixed model. F) The square root of the sum of aggressive behaviors toward the stimulus female in female-oriented aggression shown by eggs and sex. Pairs produced 1.3-fold more aggressive behaviors in the none or surrogate egg conditions relative to the own condition. Error bars show the standard error estimate from the linear mixed model that represents individual variance centered around the mean estimate from the model, *** indicates *P*< 0.001, ** indicates *P*< 0.01, * indicates *P*< 0.05, and “NS” indicates not significant.

To evaluate whether the significant difference in latency to address the stimulus animal between the own and surrogate eggs conditions might be related to variation in the age or number of eggs present in the nest between the two conditions, we analyzed latency including these variables as covariates in the linear mixed model. The average age of the eggs in the own condition was 0.44 days younger than in the surrogate conditions (Mean ± SE, Own: 2.365 ± 0.076, Surrogate ± 2.815 ± 0.084; *P*= 0.020, *t*(50) = -2.410). There were also slightly fewer eggs in the own condition than in the surrogate condition, though this difference was not significant (Mean ± SE, Own: 740.375 ± 36.189, Surrogate: 745.308 ± 48.339; *P*> 0.05). When including both terms as covariates, the effect of egg condition was no longer significant (*P*> 0.05; see Supplementary Excel File 9 for the ANOVA table including these covariates). This indicates differences between own and surrogate conditions for latency to address the stimulus male may be explained by variation in the number and age of the eggs used between the tests, rather than the fact the eggs were surrogate versus own.

Similar to the large intruder aggression assay, we found several significant effects of round. This was observed in the sum of aggressive behaviors (Square root transformed, χ^2^_2_= 13.509, *P*= 0.001, η^2^_p_= 0.082; Fig. 3B), latency (Log transformed, χ^2^_2_= 26.594, *P*< 0.001, η^2^_p_= 0.205; supplementary Fig. 1E) and displays (No transformation, χ^2^_2_= 13.765, *P*= 0.001, η^2^_p_= 0.125; supplementary Fig. 1F). Between round 1 and round 3, pairs produced 1.2 times more aggressive behaviors including 1.5 times more displays, and reduced latency over 76% to address the stimulus animal (see Supplementary Excel File 4 for pairwise comparisons, 9 for ANOVA tables, and 10 for means and standard errors).

All recorded male-oriented behaviors were significantly repeatable except for latency (see table 5 for ICC and P-values).

**Table 5.** ICC and P-value for the random effect for all measured behaviors in the male-oriented aggression assay.

| Behavioral Measure | ICC | P-Value of Random Effect |
| --- | --- | --- |
| Sum of Aggressive Behaviors | 0.220 | **<0.001** |
| Latency | 0.094 | 0.06 |
| Displays | 0.331 | **<0.001** |
| Chases/Charges | 0.275 | **<0.001** |
| Strikes | 0.300 | **<0.001** |
| Bites | 0.169 | **0.001** |

### Female-Oriented Aggression: Females are more aggressive toward intruder females than males are

For the female-oriented aggression assay, we introduced a stimulus female into the home tank of the focal pair and measured their behavioral responses. We found that in several instances, the size of the stimulus female relative to the focal female had a significant effect in our models. Thus, in all analyses of behaviors in female-oriented aggression, we added a covariate “length difference” which was the length of the focal female subtracted by the length of the stimulus female (see Supplementary Excel File 1 for lengths of animals). We also included length difference-by-sex interaction.

While we did not find a significant effect of sex on the sum of aggressive behaviors (*P*>0.05), sex differences emerged when we analyzed the behaviors separately. We observed a significant effect of sex on the number of bites (Square root transformed, χ^2^_1_= 7.846, *P*= 0.005, η^2^_p_= 0.582; Fig. 3C), strikes (No transformation, χ^2^_1_= 6.259, *P*= 0.012, η^2^_p_= 0.517; supplementary Fig. 2A), chases/charges (Square root transformed, χ^2^_1_= 4.523, *P*= 0.033, η^2^_p_= 0.252; supplementary Fig. 2B) and displays (Square root transformed, χ^2^_1_= 4.161, *P*= 0.041, η^2^_p_= 0.410; Fig. 3D). Females bit the stimulus animal over 6.5 times more than males, struck the stimulus animal 3.2 times more than males, and chased the stimulus animal 4.1 times more than males, while males produced nearly double the number of displays. Taken together, these data suggest males preferred indirect aggression (displays), while females preferred direct aggression (bites, strikes, chases/charges) (see Supplementary Excel File 11 for ANOVA tables, and 12 for means and standard errors).

The difference in length between the focal female and stimulus female had a significant effect on bites (Square root transformed, χ^2^_1_= 5.684, *P*= 0.017, η^2^_p_= 0.022; Fig. 3E), and strikes (No transformation, χ^2^_1_= 4.253, *P*= 0.039, η^2^_p_= 0.013; supplementary Fig. 2C). In both cases, pairs were less aggressive towards the stimulus female as the size of the stimulus female increased relative to the focal female. Specifically, pairs produced approximately 0.9 fewer bites and strikes per centimeter length difference (Fig. 3E and supplementary Fig. 2C). The interaction between length difference and sex was not significant (*P*> 0.05) (see Supplementary Excel File 11 for ANOVA table).

We also found a significant effect of egg condition on the sum of aggressive behaviors (Square root transformed, χ^2^_2_= 25.446, *P*< 0.001, η^2^_p_= 0.141; Fig. 3F). Evaluating behaviors individually, this effect was present for chases/charges (Square root transformed, χ^2^_2_= 14.743, *P*< 0.001, η^2^_p_= 0.101; supplementary Fig. 2D), displays (Square root transformed, χ^2^_2_= 9.682, *P*= 0.008, η^2^_p_= 0.057; supplementary Fig. 2E), and strikes (No transformation, χ^2^_2_= 10.462, *P*= 0.005, η^2^_p_= 0.066; supplementary Fig. 2F). Similar to male-oriented aggression, pairs tended to be more aggressive toward the stimulus female in the none or surrogate conditions compared to the own condition. Relative to the own condition, pairs in the none or surrogate conditions displayed 1.3-fold more aggressive behaviors, 2.2-fold more chases/charges, 1.5-fold more strikes, and 1.1-fold more displays. For all the above measures, no significant difference was detected between the none and surrogate eggs conditions (see supplementary Excel File 4 for pairwise comparisons, 11 for ANOVA tables, and 12 for means and standard errors).

Similar to what was described in male-oriented aggression, to evaluate whether increased female-oriented aggression in the surrogate condition relative to the own condition could be related to variation in the age or number of eggs between the tests, we again included these variables as covariates in analyses including only own or surrogate egg conditions. The age of eggs had a significant negative effect on the sum of aggressive behaviors (Square root transformed, χ^2^_1_= 10.101, *P*= 0.001, η^2^_p_= 0.113; see supplementary Fig. 2G) and displays (Square root transformed, χ^2^_1_= 5.919, *P*= 0.015, η^2^_p_= 0.068; see supplementary Fig. 3H) but was not significant for strikes or chases/charges (*P*> 0.05). Pairs exhibited approximately 6 fewer total aggressive behaviors and 2 fewer displays as the age of the eggs increased per day. The count of eggs also had a significant negative effect on the sum of aggressive behaviors (Square root transformed, χ^2^_1_= 3.926, *P*= 0.048, η^2^_p_= 0.046; supplementary Fig. 3I). Pairs exhibited approximately 0.6 fewer aggressive behaviors per 100 eggs in the nest. However, even when including these covariates in the mixed model, the difference between own and surrogate conditions remained significant for all of the aforementioned variables (see Supplementary Excel File 11 for ANOVA tables including these covariates). This suggests that there may have been some subtle differences between the own versus surrogate conditions that cannot be explained by variation in the number or age of the eggs during the tests (see Discussion).

There was also a significant interaction between egg condition and sex for the sum of aggressive behaviors (Square root transformed, χ^2^_2_= 6.587, *P*= 0.037, η^2^_p_= 0.047; supplementary Fig. 3A). Females displayed 37% fewer aggressive behaviors in the own condition compared to the none and surrogate conditions, while males showed no significant difference in aggression based on egg condition. This interaction remained significant when including the age and count of eggs as covariates (see Supplementary Excel File 11 for the ANOVA table including these covariates).

Similar to the large intruder aggression and male-oriented aggression assays, we again found a significant effect of round on the sum of aggressive behaviors (Square root transformed, χ^2^_2_= 8.173, *P*= 0.017, η^2^_p_= 0.120; supplementary Fig. 3B), displays (Square root transformed, χ^2^_2_= 6.125, *P*= 0.047, η^2^_p_= 0.145; supplementary Fig. 3C), strikes (No transformation, χ^2^_2_= 7.899, *P*= 0.019, η^2^_p_= 0.036; supplementary Fig. 3D), and latency (Log transformed, χ^2^_2_= 22.238, *P*< 0.001, η^2^_p_= 0.187; supplementary Fig. 3E). Pairs got more aggressive, and more rapidly addressed the stimulus female as rounds progressed. The total number of aggressive behaviors increased 1.4-fold between round 1 and 3. Broken down by individual behaviors, we observed a 1.6-fold increase in displays, a 1.1-fold increase in strikes, and a 60% decrease in latency to address the stimulus female between round 1 and round 3 (see Supplementary Excel File 4 for pairwise comparisons, 11 for ANOVA tables, and 12 for means and standard errors).

All measured female-oriented behaviors were significantly repeatable except latency (see Table 6 for ICC and P-values).

**Table 6.** ICC and P-value for the random effect for all measured behaviors in the female-oriented aggression assay.

| Behavioral Measure | ICC | P-Value of Random Effect |
| --- | --- | --- |
| Sum of Aggressive Behaviors | 0.403 | **<0.001** |
| Latency | 0.069 | 0.137 |
| Displays | 0.273 | **<0.001** |
| Chases/Charges | 0.238 | **<0.001** |
| Strikes | 0.250 | **<0.001** |
| Bites | 0.339 | **<0.001** |

### Aggression Ratio: Females have a higher ratio than males

We replicated the finding in (Parker et al., 2022) that females displayed approximately 2.7 fold higher aggression ratio than males (sum of aggressive behaviors in the female-oriented aggression assay divided by the sum of aggressive behaviors in the male-oriented aggression assay; Log10 transformed, χ^2^_1_= 6.881, *P*= 0.009, η^2^_p_= 0.392; Fig. 4A). No significant effects of egg condition, round, or any of the interactions were detected (*P* >0.05; see Supplementary Excel File 13 for the ANOVA table, and 14 for means and standard errors).

**Figure 4.**
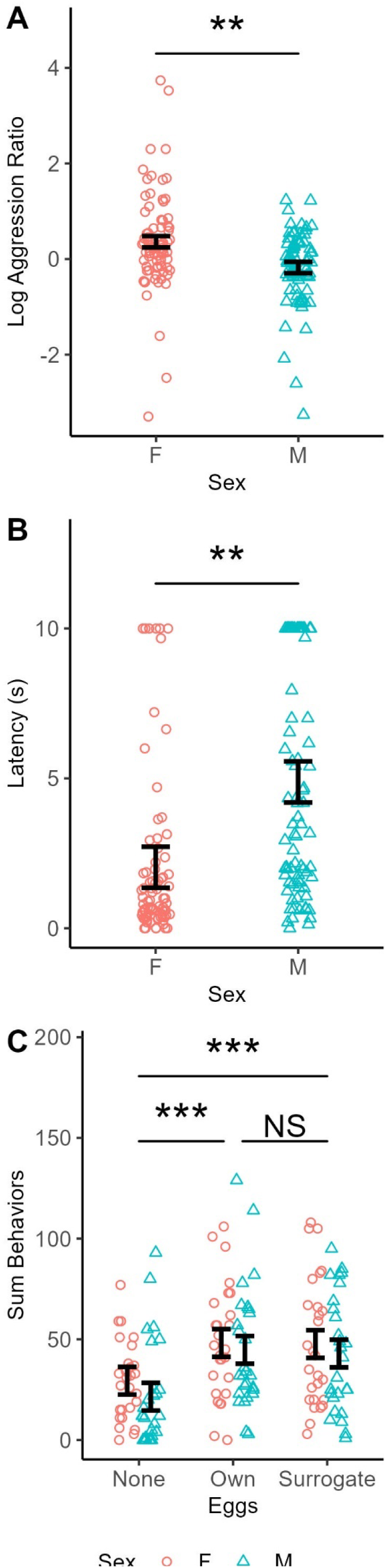
Females are bolder than males and nest maintenance increases when eggs are in the nest. All datapoints are shown including repeated measures. A) Log10 aggression ratio shown by sex. Females are 2.7-fold more biased toward female-oriented aggression than males. B) Latency to address the stimulus in immediate reaction to a threat shown by sex. Females were 35% faster to respond to the stimulus than males. C) The sum of behaviors aimed at removing the contaminant in nest maintenance shown by egg condition and sex. Pairs devoted more effort to remove the contaminant with eggs in the nest compared to without. Error bars show the standard error estimate from the linear mixed model that represents individual variance centered around the mean estimate from the model, *** indicates *P*< 0.001, ** indicates *P*< 0.01, and “NS” indicates not significant.

The female-male aggression ratio was not significantly repeatable (Log transformed, ICC: 0.058, Random effect *P*= 0.214) suggesting within individual variation was magnified by taking the ratio of behaviors across two assays.

### Immediate Reaction to a Threat: Females are significantly bolder than males

To measure boldness, we measured the focal animal’s immediate response to a threat (a researcher’s hand) within a short timeframe (10s). We found a significant effect of sex on the number of chases/charges (Log transformed, χ^2^_1_= 4.952, *P*= 0.026, η^2^_p_= 0.244; supplementary Fig. 4A). In fact, no males were observed charging whereas 4 females performed this behavior. Females were also 59% faster to respond to the stimulus than males (No transformation, χ^2^_1_= 7.782, *P*= 0.005, η^2^_p_= 0.350; Fig. 4A; see Supplementary Excel File 15 for ANOVA tables, and Excel File 16 for means and standard errors).

While there were no significant main effects of sex or egg condition for displays, there was a significant interaction between these factors (No transformation, χ^2^_2_= 6.319, *P*= 0.042, η^2^_p_= 0.044; supplementary Fig. 4B), though no pairwise contrasts were significant. We also found a significant interaction between sex and egg condition for latency (No transformation, χ^2^_2_= 10.063, *P*= 0.007, η^2^_p_= 0.069; supplementary Fig. 4C). Post hoc tests indicated that females did not significantly differ in latency based on egg condition, while males addressed the stimulus 33% faster with no or surrogate eggs in the nest relative to own eggs (see Supplementary Excel File 4 for pairwise comparisons, 15 for ANOVA tables, and 16 for means and standard errors).

To evaluate whether the difference in latency to address the stimulus in males between the own and surrogate conditions could be related to variation in the age or number of eggs between the tests, we again included these as covariates in an analysis of only these two egg conditions. Neither of these covariates were significant, and the significant difference between own and surrogate egg conditions remained (see Supplementary Excel file 15 for the ANOVA table including these covariates). This suggests the slight delay that males have in addressing an immediate threat when caring for their own versus surrogate eggs cannot be explained by variation in the number or age of the eggs between the tests.

All recorded behaviors were significantly repeatable except for number of strikes and bites (see table 7 for ICC and P-values). This is likely caused by the rarity of strikes and bites recorded in this assay. Across the 81 videos analyzed, there were a total of 8 bites and 8 strikes recorded, with 7 of the bites and 7 of the strikes produced by 3 females (C11F, C15F, and T18F).

**Table 7.** ICC and P-value for the random effect for all measured behaviors in the immediate reaction to a threat assay.

| Behavioral Measure | ICC | P-Value of Random Effect |
| --- | --- | --- |
| Sum of Aggressive Behaviors | 0.471 | **<0.001** |
| Latency | 0.322 | **<0.001** |
| Displays | 0.385 | **<0.001** |
| Chases/Charges | 0.251 | **<0.001** |
| Strikes | 0.092 | 0.058 |
| Bites | 0.069 | 0.142 |

### Nest Maintenance

To assess if there was a division of labor in nest maintenance, we introduced a nest contaminant (a small rock) into the terra cotta pot, and measured behaviors aimed at removing the contaminant from the nest (strikes, bites, and latency). We found a slight but significant effect of sex on the sum of aggressive behaviors (No transformation, χ^2^_1_= 4.081, *P*= 0.043, η^2^_p_= 0.025; supplementary Fig. 4D) and latency to address the stimulus (Log transformed, χ^2^_1_= 5.943, *P*= 0.015, η^2^_p_= 0.005; supplementary Fig. 4E). Females performed 1.1-fold more aggressive behaviors than males and addressed the contaminant 35% faster than males (see Supplementary Excel File 17 for ANOVA tables, and 18 for means and standard errors).

We also found a significant effect of egg condition on a several recorded behaviors. This included the sum of behaviors (No transformation χ^2^_2_= 12.246, *P*= 0.002, η^2^_p_= 0.180; Fig. 4B), strikes (No transformation, χ^2^_2_= 10.809, *P*= 0.004, η^2^_p_= 0.198; supplementary Fig. 4F), and bites (Square root transformation, χ^2^_2_= 9.436, *P*= 0.009, η^2^_p_= 0.115; supplementary Fig. 5A). Pairs were significantly more aggressive toward the contaminant with eggs in the nest (own or surrogate) as opposed to without. No differences between own and surrogate egg conditions were detected. Pairs responded to the contaminant 63% faster with eggs in the nest compared to without and produced nearly double the number behaviors aimed at removing the contaminant (see Supplementary Excel File 4 for pairwise comparisons, 17 for ANOVA tables, and 18 for means and standard errors).

In addition, we found a significant effect of round on strikes (No transformation, χ^2^_2_= 6.883, *P*= 0.032, η^2^_p_= 0.003; supplementary Fig. 5B). Similar to previous assays, as rounds progressed, pairs got more aggressive, though no contrasts were significant in post hoc tests (see Supplementary Excel File 4 for pairwise comparisons, 17 for ANOVA tables, and 18 for means and standard errors).

We also found a significant interaction between sex and round on the sum of behaviors (No transformation, χ^2^_2_= 8.399, *P*= 0.015, η^2^_p_= 0.058; supplementary Fig. 5C), strikes (No transformation, χ^2^_2_= 10.525 *P*= 0.005, η^2^_p_= 0.072; supplementary Fig. 5D), and latency (Log transformed, χ^2^_2_= 10.761, *P*= 0.005, η^2^_p_= 0.073; supplementary Fig. 5E). Males significantly increased nest maintenance behaviors and decreased latency as rounds progressed, while females significantly decreased strikes as rounds progressed (see Supplementary Excel File 4 for pairwise comparisons, 17 for ANOVA tables, and 18 for means and standard errors).

Further, we found a significant interaction between sex and eggs for latency (Log transformed, χ^2^_2_= 10.761, *P*= 0.004, η^2^_p_= 0.073; supplementary Fig. 5F). Males addressed the contaminant significantly faster with eggs in the nest compared to without, while females did not significantly differ based on egg condition (see Supplementary Excel File 4 for pairwise comparisons, 17 for ANOVA tables, and 18 for means and standard errors).

All recorded nest maintenance behaviors were significantly repeatable (see table 8 for ICC and P-values).

**Table 8.** ICC and P-value for the random effect for all measured behaviors in the nest maintenance assay.

| Behavioral Measures | ICC | P-Value of Random Effect |
| --- | --- | --- |
| Sum of Behaviors | 0.340 | **<0.001** |
| Latency | 0.375 | **<0.001** |
| Strikes | 0.500 | **<0.001** |
| Bites | 0.509 | **<0.001** |

### Principal Component Analysis (PCA)

In addition to analyzing each behavioral assay separately to determine effects of sex, round and egg condition, we also sought to evaluate how a composite of all of the recorded assays might be a useful metric for differentiating the sexes. Thus, we performed a PCA analysis on the sum of behaviors from each assay collapsed across all rounds and egg conditions within individuals. We found that the first and second PCs explained 65.7% of the variation. A plot of PC2 on PC1 shows the individuals largely separate based on a parental-aggression axis (Fig. 5) which effectively separates the sexes, except for a few individuals at the margins. These include 1 male (T18M in Fig. 5A) which displays similar levels of aggression and parental care as females, and 2 females that do more parenting and display less aggression than usual (A24F and B22F in Fig. 5A). PC1 from this composite PCA analysis significantly differentiated the sexes (*t*(16)= 3.512, *P*= 0.003), whereas PC2 did not (*P*>0.05). Additionally, we found that PC2 captured the variation among aggression assays (Fig. 5B). Conspecific-oriented aggression assays loaded negatively, while non-conspecific aggression assays loaded positively (with the exception of immediate reaction to a threat), suggesting conspecific aggression and non-conspecific aggression constitute two distinct types of aggression.

**Figure 5.**
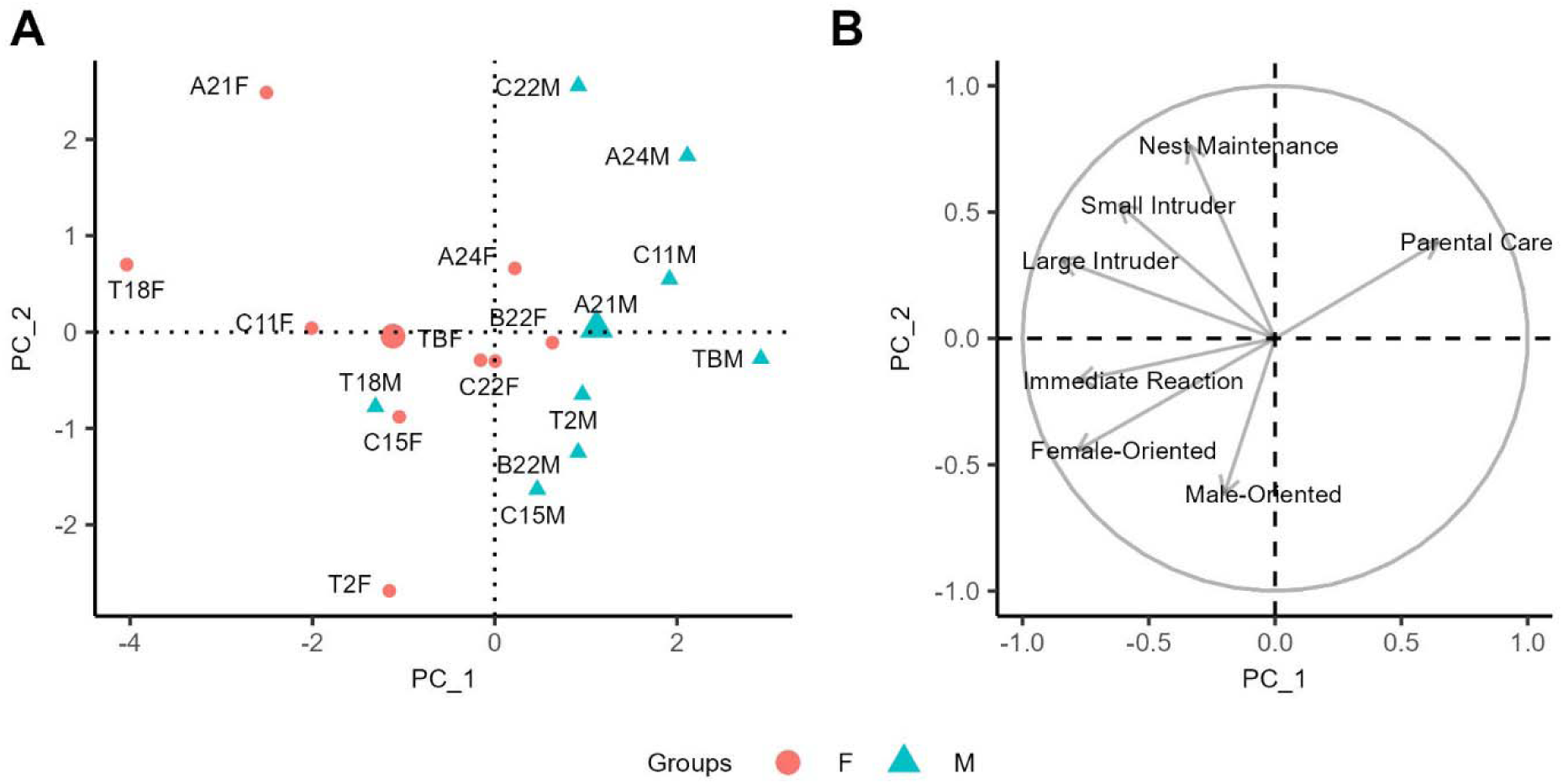
Males and females separate largely along a parental care - aggression axis. A) A PCA was performed on the sum of behaviors (collapsed across rounds and egg conditions) for each of the 7 behavioral assays. PC1 is plotted against PC2 showing males and females separately. In addition, the global female and male averages are shown as large red circle and large blue triangle respectively. B) Loadings of the corresponding assays are shown on PC1 and PC2. Males and females largely split along a parental care - aggression axis, where parental care loads to quadrant 1 while the various aggression assays load to quadrants 2 and 3.

Additionally, we performed multiple separate PCA analyses for all of the recorded behaviors within each assay (supplementary Figs. 6-8). We then calculated the Euclidean distance between the mean male and mean female PC1 and PC2 coordinates. Based on this metric, the assays that best differentiated the sexes were parental care (2.767), female-oriented aggression (2.645), and large intruder aggression (2.297; table 9). PC1 extracted from the analyses of parental care, large intruder aggression, female-oriented aggression, and immediate reaction to a threat significantly differentiated males and females but did not significantly differentiate the sexes in the other assays, while PC2 extracted from male-oriented aggression significantly differentiated the sexes (see Supplementary Excel File 19 for statistical analyses). When analyzing each assay separately, a few consistent patterns emerged. As in our statistical analysis, in parental care, males are characterized by high levels of parental behaviors and time spent in the nest (supplementary Figs. 6A, 6B). In non-conspecific aggression assays (large intruder, small intruder, immediate reaction, nest maintenance), males differentiate from females based on high latency and low direct aggression (bites, strikes, chases/charges) (supplementary Figs. 6,8). Similarly, in conspecific-oriented aggression assays, males differentiate from females based on direct versus indirect aggression, with males characterized by high indirect aggression, and females characterized by high direct aggression, while latency does not differentiate the sexes (supplementary Fig. 7).

**Table 9.** Euclidian distances between mean male and mean female PC1 and PC2 coordinates. Parental care, female-oriented aggression, and large intruder aggression produce the largest Euclidean distance between the average male and average female.

| Assay | Distance |
| --- | --- |
| Parental Care | 2.767 |
| Small Intruder Aggression | 1.165 |
| Large Intruder Aggression | 2.297 |
| Male-Oriented Aggression | 1.565 |
| Female-Oriented Aggression | 2.645 |
| Immediate Reaction to a Threat | 2.147 |
| Nest Maintenance | 0.577 |

### Pair Dynamics

To determine whether the male-female pairs coordinated their behaviors toward the introduced stimuli, we estimated Pearson’s correlations separately for each assay using the sum of behaviors collapsed across round and egg conditions. The only correlation that was significant was female-oriented aggression where levels of aggression in the male and female of the pair were positively correlated (*r*(7)= 0.720, *P*= 0.029; supplementary Fig. 9).

### Behavioral Syndromes

To assess the presence of behavioral syndromes, we fit Pearson’s correlations in a pairwise manner between all 7 behavioral assays. For this analysis, we used the mean of sum of behaviors collapsed across round and egg condition to represent individual values for each assay. This resulted in a total of 21 comparisons. Because we observed several significant effects of sex, we assessed behavioral syndromes for males and females separately.

#### Males

Of the 21 correlations calculated for males, 5 were significant. Large intruder aggression was significantly positively correlated with small intruder aggression (*r*(7)= 0.710, *P*= 0.032; Fig. 6A). Both female-oriented aggression and male-oriented aggression were significantly positively correlated with immediate reaction to a threat (*r*(7)= 0.897, *P*= 0.001; Fig. 6B; *r*(7)= 0.831, *P*= 0.006; Fig. 6C, respectively). Female-oriented aggression was also positively correlated with male-oriented aggression (*r*(7)= 0.819, *P*= 0.007; Fig. 6D), and negatively correlated with parental care (*r*(7)= -0.688, *P*= 0.041; Fig. 6E).

**Figure 6.**
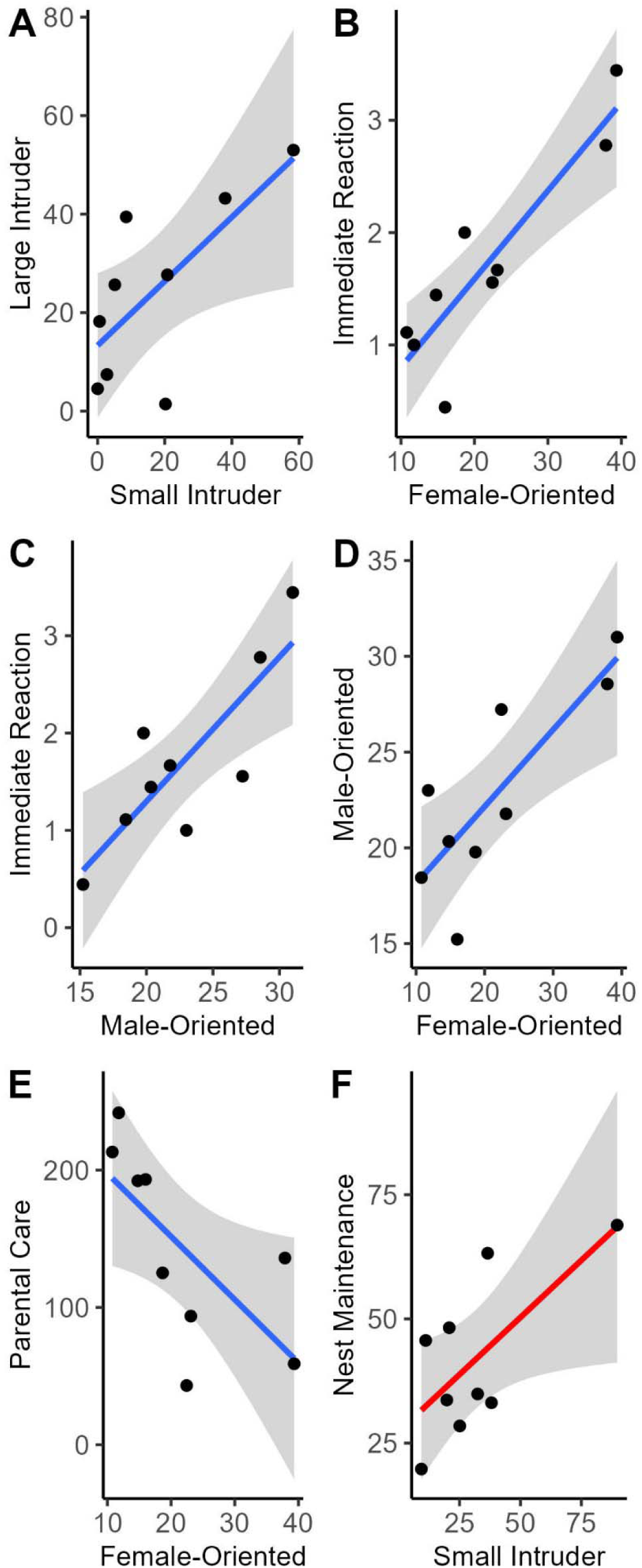
Males display several behavioral syndromes, while females display only one. All data points represent the sum of behaviors collapsed across round and egg condition within each individual A) The sum of aggressive behaviors in large intruder aggression plotted against the sum of aggressive behaviors in small intruder aggression, males only (*P*= 0.032). B) The sum of aggressive behaviors in immediate reaction to a threat plotted against the sum of aggressive behaviors in female-oriented aggression assay, males only (*P*= 0.001). C) The sum of aggressive behaviors in immediate reaction to a threat plotted against the sum of aggressive behaviors in male-oriented aggression, males only (*P*= 0.006). D) The sum of aggressive behaviors in male-oriented aggression plotted against sum of aggressive behaviors in female-oriented aggression, males only (*P*= 0.007). E) The sum of parental care behaviors) plotted against the sum of aggressive behaviors in female-oriented aggression, males only (*P*= 0.042). F) The sum of behaviors in nest maintenance assay plotted against the sum of aggressive behaviors in small intruder aggression, females only (*P*= 0.041). Blue and red lines represent simple linear regression lines for males and females, respectively. Ribbons show ± 95% confidence interval.

#### Females

Surprisingly, of the 21 correlations we calculated for females, only 1 was significant. Small intruder aggression was significantly correlated with nest maintenance (*r*(7)= 0.685, *P*= 0.042; Fig. 6F). This is perhaps unsurprising due to the high visual similarity between a hermit crab in the small intruder aggression assay and a rock in the nest maintenance assay, though it is unclear why this syndrome is detectable in females but not males.

## Discussion

Here we significantly broadened the range of quantifiable behaviors in *A. ocellaris*, enabling a more comprehensive examination of their unique characteristics in a laboratory setting. To this end, we established significant individual differences, effects of sex, effects of the environment (e.g., whether eggs are present or not), and effects of repeated testing (e.g., round) on multiple measures of territory defense, aggression, and parental care. All assays contained measures that were significantly repeatable across contexts and rounds. The average ICC of measured behaviors was 0.33, consistent with average estimates of repeatability of behavior (Bell et al., 2009). Interestingly, in male and female-oriented aggression, latency was not significantly repeatable. This lack of repeatability could be due to a significant effect of the behavior of the stimulus animal, or spontaneity in the decisions to engage. The female-male aggression ratio was also not significantly repeatable which is likely due to its compound nature combining two partially repeatable behaviors: male and female-oriented aggression (Certo et al., 2020; Jasieński & Bazzaz, 1999).

All of the assays described in this paper except for male-oriented aggression had at least one behavior that significantly differed based on sex. This was almost always an increase in aggression in females, and an increase in latency in males. Interestingly, in female-oriented aggression, the sexes did not significantly differ in the sum of aggressive behaviors, but did differ in their behavioral composition, with males displaying indirect aggression (displays) while females displaying more direct aggression (bites) (Figure 3C, 3D). Thus, future studies that use this assay to evaluate sex differences should evaluate displays and bites separately. The different ways in which males and females direct aggression towards a female intruder are likely due to the significant size difference between males and females and their unique natural history. Males may be less willing to directly confront a female because of possible injury, whereas a female will fight to the death to maintain / usurp the α position in the hierarchy as searching for a new anemone means almost certain death (Da Silva & Nedosyko, 2016; Iwata & Manbo, 2013).

We found that the best assays for detecting sex differences were parental care, female-oriented aggression, and large intruder aggression. These assays detected large, and significant sex effects across environmental conditions and rounds. Because of the large effect size and internal consistency, these assays would be the most useful for evaluating the behavioral sex of a fish in the midst of sex change. However, we found that none of these, whether considered separately or combined into a linear combination could perfectly distinguish the sexes. This was due to strong and repeatable individual differences in behavior with some females behaving similarly to males (e.g., B22F in Fig. 5A and supplementary Figs. 6C, 7A, 7C, and 8A), and some males behaving similarly to females (e.g., T18M in Fig. 5A and supplementary Figs. 6A, 6C) in specific assays.

In addition to uncovering large sex differences for independent, repeatable behaviors, we also observed strong evidence for sex specific behavioral syndromes in *A. ocellaris*. In males, we observed positive correlations between several aggression measures and boldness, and a negative correlation between aggression and parental care, as shown earlier (R. DeAngelis et al., 2017, 2020). However, in females, only one correlation was observed for aggression toward a rock (nest maintenance) and a hermit crab (small intruder). Most of the sex differences we observed including the syndromes are likely caused at some level by the organizational and activational effects of the sex steroid hormones 11-ketotestosterone (11-KT) and estradiol (E2), which have been shown to have broad effects on multiple aspects of physiology and behavior (Gegenhuber et al., 2022; Godwin, 2010; Stennette & Godwin, 2024). Additionally, it may be the case that females are more variable within sex than males, resulting in our statistical analyses not being able to detect behavioral syndromes in females given the small sample size.

In anemonefish, male-female pairs work together as a reproductive unit (Laudet & Ravasi, 2022), yet our study shows that an individual’s behavior was largely independent of the behavior of its partner, with the exception of female-oriented aggression. This is likely because of the strong division of labor, where individuals perform their duties independently without much help or compensation from the other (R. DeAngelis et al., 2017). The correlation between male and female aggression levels in the female-oriented aggression assay could reflect the size differential between males and females. A male is probably unwilling to be aggressive towards a much larger female unless its female partner is also engaging with him for fear of retaliation.

In many of our assays, we found at least one behavior that significantly changed as rounds progressed. This was almost always an increase in aggressive behavior or a decrease in latency to address the stimulus. These round differences are likely caused by acclimation to the stimulus following repeated exposure leading to increased confidence that aggression does not pose a risk. In male and female-oriented aggression, the increase of aggression as round progressed may also be caused by winner effects as, in most cases, the focal pair would likely perceive they “won” the encounter (Dijkstra et al., 2012; Dugatkin, 1997; Hsu & Wolf, 1999).

We also found in many assays that the fish’s behavior depended on the context, whether pairs had their own, surrogate, or no eggs present in the nest when the behaviors were recorded. In nest maintenance, when the pairs had no eggs in the nest, they displayed fewer behaviors aimed at clearing the contaminant than when they had their own or surrogate eggs. This was expected, as a contaminant in the nest serves as an obstacle to care of the eggs meaning motivation to clear the contaminant would be higher with eggs in the nest compared to without. Additionally, when eggs are in the nest, the male spends more time in the nest than when it is empty (R. S. DeAngelis & Rhodes, 2016), meaning the male is more likely to notice the contaminant, and will more rapidly address it (supplementary Fig. 5F). However, we did not expect to find the consistent differences between own and surrogate conditions we observed. There were significant differences in behavior between the own and surrogate conditions in male-oriented aggression (Fig. 3A), female-oriented aggression (Fig. 3F, supplementary Figs. 2E, 2F, 3A), and immediate reaction to a threat (supplementary Fig. 4C) that were consistently characterized by increased latency and decreased aggression in the own condition relative to none or surrogate. For the most part, we were not able to explain these significant differences with the count or age of the eggs suggesting two possibilities. The first possibility is that pairs can differentiate their own eggs from surrogate eggs, though this seems unlikely as own and surrogate eggs are treated the same in all other assays. The second possibility is that some characteristic of the experiment that could not be accounted for lead to subtle changes in behavior in these assays. Replacing the terra cotta pot serving as the pair’s nest with an unfamiliar terra cotta pot in the surrogate condition could explain this difference. Even after the 24h acclimation period in the surrogate condition, it may be the case that the act of switching pots leads to an increase in stress leading to more aggression in the surrogate condition than in the own condition. Future studies should examine under what contexts pairs can detect differences between own and surrogate eggs to better understand the scope of this difference.

In conclusion, we measured the behavior of *A. ocellaris* pairs in 6 novel behavioral assays and 1 established parental care assay. All behaviors were measured across 3 contexts that varied depending on whether no eggs, own or surrogate eggs were present. In addition, all behaviors were measured 3 separate times during separate spawning cycles to evaluate repeatability. Thirty-four out of 39 behaviors showed significant repeatability. The best assays for differentiating the sexes were the parental care assay, large intruder aggression assay, and female-oriented aggression assay. Additionally, we found consistent differences in aggression depending on egg condition and level of repetition, where pairs tended to display reduced aggression with their own eggs relative to surrogate or no eggs, and increased aggression with repeated exposure to the stimulus. Further, we found that the behaviors of males and females within a pair were largely uncorrelated, except for female-oriented aggression. Interestingly, males displayed several behavioral syndromes, but females only displayed one. This study significantly broadens the number of laboratory behavioral assays available for studying female behavioral dominance and territoriality, protandrous behavioral sex change, behavioral plasticity and the neuroendocrine basis of parental care and aggression in the iconic anemonefish, *A. ocellaris*.

