## supplementary Fig. for "Novel Behavioral Assays Reveal Sex Specific Behavioral Syndromes in Anemonefish"

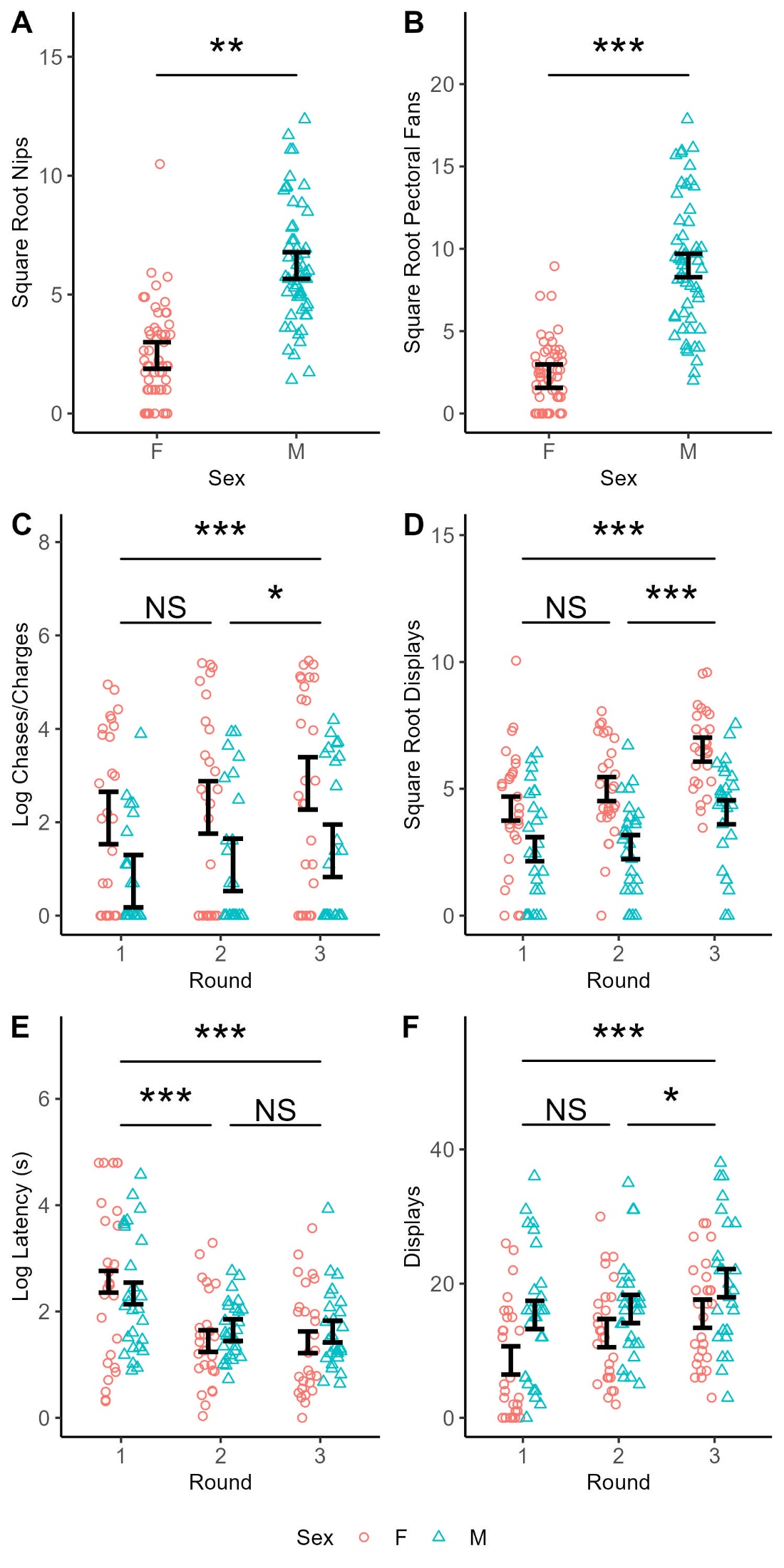

**Supplementary Figure 1. Males perform more parental care behaviors and pairs get more aggressive as rounds progress.** All data points are shown including repeated measures. A) The square root of nips in parental care shown by sex. Males performed 4.6-fold more nips than females. B) The square root pectoral fans in parental care shown by sex. Males performed 10.2-fold more pectoral fans than females. C) The log of chases/charges toward the stimulus animal in large intruder aggression shown by round and sex. Pairs performed 2.5-fold more chases/charges as rounds progressed. D) The square root displays in large intruder aggression shown by sex and round. Pairs produced 1.9-fold more displays as rounds progress. E) The log of latency to address the stimulus male in male-oriented aggression shown by round and sex. Pairs addressed the stimulus male 76% faster as rounds progressed. F) Displays in male-oriented aggression shown by round and sex. Pairs produced 1.5-fold more displays as rounds progress. Error bars show the standard error estimate from the linear mixed model that represents individual variance centered around the mean estimate from the model, *** indicates *P*< 0.001, ** indicates *P*< 0.01, * indicates *P*< 0.05, and “NS” indicates not significant.

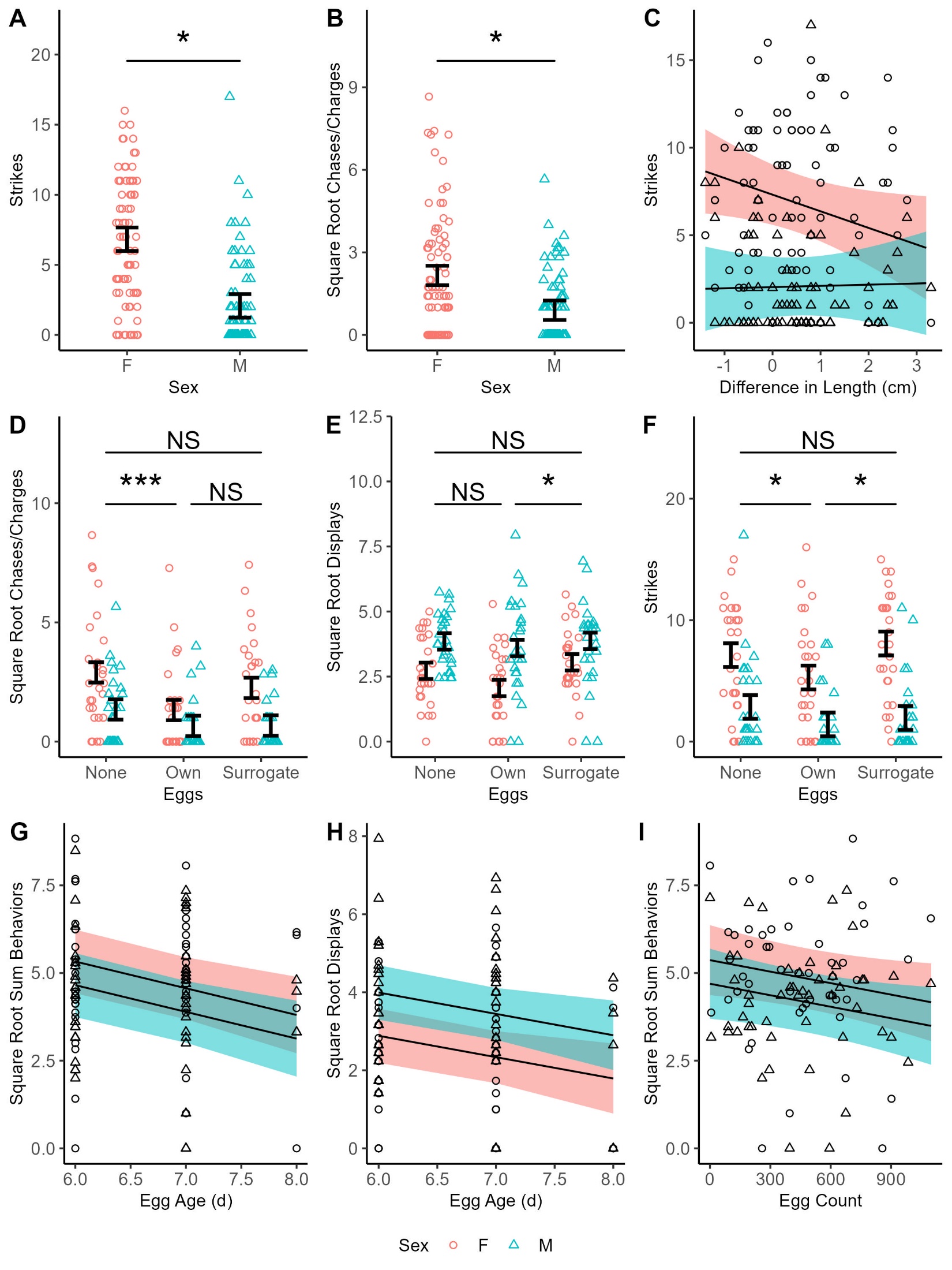

**Supplementary Figure 2. Females perform significantly more direct aggression than males, pairs get less aggressive as the focal female increases in size relative to the stimulus female, pairs are the least aggressive in the own eggs condition, and egg age and count are negative predictors of aggression.** All data points are shown including repeated measures. A) Strikes toward the stimulus female in female-oriented aggression shown by sex. Females produce 3.2-fold more strikes than males. B) The square root of chases/charges in female-oriented aggression shown by sex. Females chased the stimulus female 4.1-fold more than males. C) Strikes toward the stimulus female in female-oriented aggression shown by the difference in length between the focal female and the stimulus female and sex. Pairs got significantly less aggressive as stimulus female got larger, particularly females. D) The square root of chases/charges toward the stimulus female in female-oriented aggression shown by sex and egg condition. Pairs produced 2.7-fold more aggressive behaviors in the none condition relative to the own condition. E) The square root of displays toward the stimulus female in female-oriented aggression shown by sex and eggs. F) Strikes toward the stimulus female in female-oriented aggression shown by sex and eggs. Pairs were 32% less aggressive with own eggs relative to none or surrogate. G) The square root of the sum of aggressive behaviors in female-oriented aggression shown by egg age and sex. For each day the age of the eggs increased, the sum of aggressive behaviors decreased by 6. H) The square root of the sum of displays in female-oriented aggression shown by egg age and sex. For each day the age of the eggs increased, the number of displays decreased by 2. I) The square root of the sum of behaviors in female-oriented aggression shown by egg count and sex. For each 100 egg increase, the sum of aggressive behaviors decreased by 0.6. In C and G-I, lines represent linear regression estimates and ribbons represent the ± 95% confidence interval from the mixed model. Error bars show the standard error estimate from the linear mixed model that represents individual variance centered around the mean estimate from the model, *** indicates *P*< 0.001, * indicates *P*< 0.05, and “NS” indicates not significant.

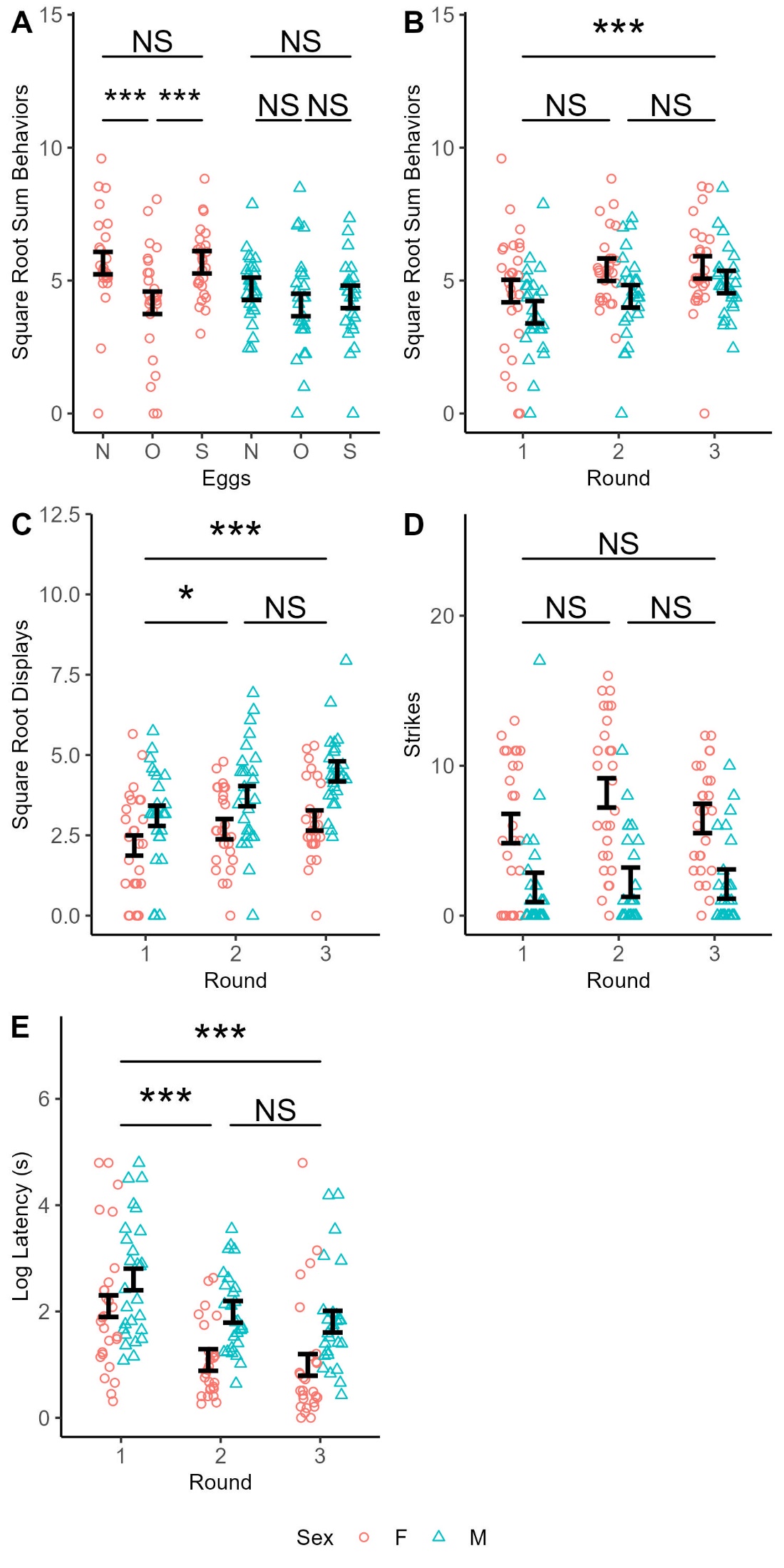

**Supplementary Figure 3. Females significantly differ in aggression based on egg condition and pairs get more aggressive as rounds progress.** All data points are shown including repeated measures. A) The square root of the sum of aggressive behaviors in female-oriented aggression shown as an eggs by sex interaction. ‘N’ indicates no eggs, ‘O’ indicates own eggs, and ‘S’ indicates surrogate eggs. Females significantly reduced aggression in the own eggs condition compared to the none or surrogate conditions, while males did not. B) The square root of the sum of aggressive behaviors in female-oriented aggression shown by sex and round. Pairs produced 1.4-fold more aggressive behaviors between round 1 and round 3. C) The square root of displays in female-oriented aggression shown by sex and round. Pairs produced 1.7-fold more aggressive behaviors between round 1 and round 3.D) Strikes in female-oriented aggression shown by sex and round. While this term was significant in the ANOVA, no pairwise comparisons were significant. E) The log of latency to address the stimulus female in female-oriented aggression shown by sex and round. Pairs addressed the stimulus female 59% faster between round 1 and round 3. Pairs produced 1.2-fold more displays in the surrogate eggs condition compared to the own eggs condition. Error bars show the standard error estimate from the linear mixed model that represents individual variance centered around the mean estimate from the model, *** indicates *P*< 0.001, * indicates *P*< 0.05, and “NS” indicates not significant.

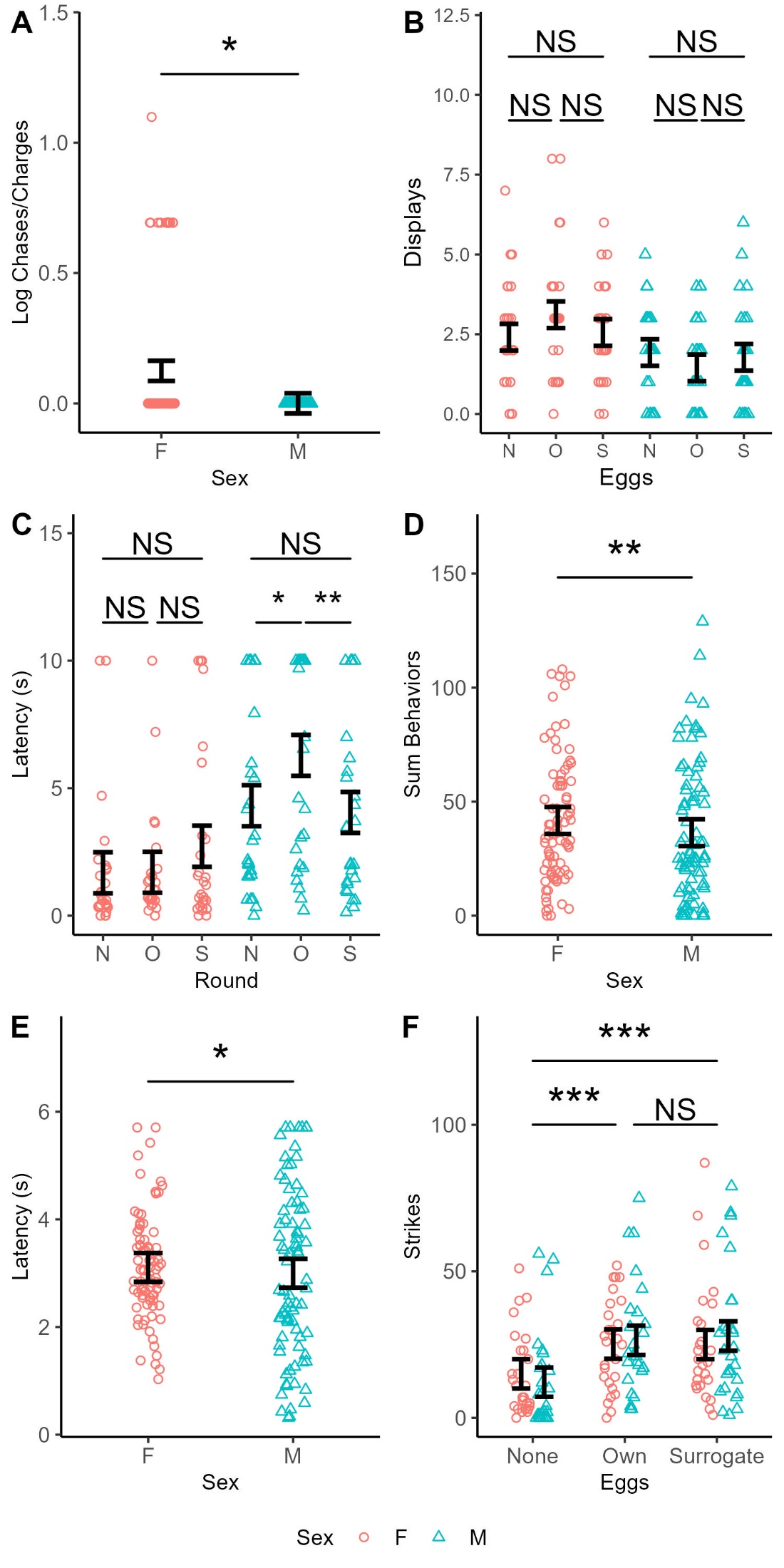

**Supplementary Figure 4. Males are less bold than females, and significantly differ in boldness based on egg condition, and females perform more nest maintenance behaviors than males. Nest maintenance behaviors are dependent on presence of eggs in the nest.** All data points are shown including repeated measures. A) The log of chases/charges in immediate reaction to a threat shown by sex. Only females were observed performing chases/charges. B) Displays toward the stimulus in immediate reaction to a threat shown as a sex by eggs interaction. While this term was significant in the ANOVA, no pairwise comparisons were significant. C) Latency to address the stimulus in immediate reaction to a threat shown as a sex by eggs interaction. While this term was significant in the ANOVA, no pairwise comparisons were significant. D) The sum of behaviors in nest maintenance shown by sex. Females performed 1.1-fold more behaviors than males. E) Latency in nest maintenance shown by sex. Females addressed the contaminant 33% faster than females. F) Strikes in nest maintenance shown by sex and eggs. Pairs produced 1.9-fold more strikes with eggs in the nest compared to without. Error bars show the standard error estimate from the linear mixed model that represents individual variance centered around the mean estimate from the model, ** indicates *P*< 0.01, * indicates *P*< 0.05, and “NS” indicates not significant.

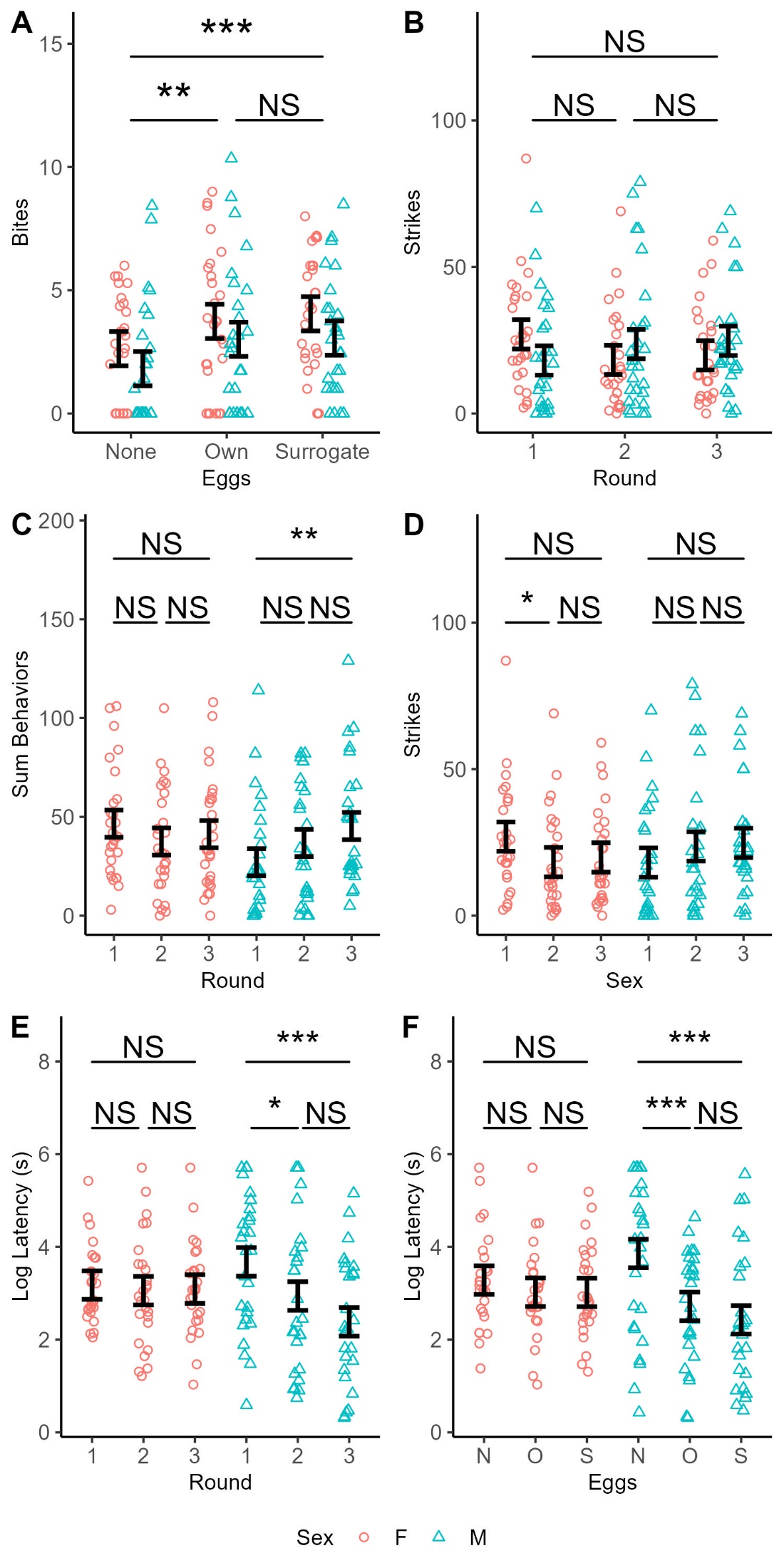

**Supplementary Figure 5. Pairs perform less nest maintenance behaviors without eggs in the nest and differences in behavior based on round or egg condition are largely driven by males.** All data points are shown including repeated measures. A) Bites toward the contaminant in nest maintenance shown by sex and eggs. Pairs produced 1.9-fold more bites with eggs in the nest compared to without. B) Strikes toward the contaminant in nest maintenance shown by sex and round. While this term was significant in the ANOVA, no pairwise comparisons were significant. C) The sum of behaviors in nest maintenance shown as a sex by round interaction. Females did not significantly differ as rounds progressed, while males performed 1.7-fold more aggressive behaviors between round 1 and round 3. D) Strikes in nest maintenance shown as a sex by round interaction. Males did not significantly change as rounds progressed, while females produced 20% fewer strikes between round 1 and round 2. E) The log of latency to address the contaminant in nest maintenance shown as a sex by round interaction. Females did not significantly change in latency to address the contaminant as rounds progressed, while males addressed the contaminant 72% faster in round 3 compared to round 1. F) The log of latency in nest maintenance shown as a sex by eggs interaction. Females did not significantly differ in latency to address the contaminant based on egg condition, while males addressed the contaminant 75% faster with eggs in the nest compared to without. Error bars show the standard error estimate from the linear mixed model that represents individual variance centered around the mean estimate from the model, *** indicates *P*< 0.001, ** indicates *P*< 0.01, * indicates *P*< 0.05, and “NS” indicates not significant.

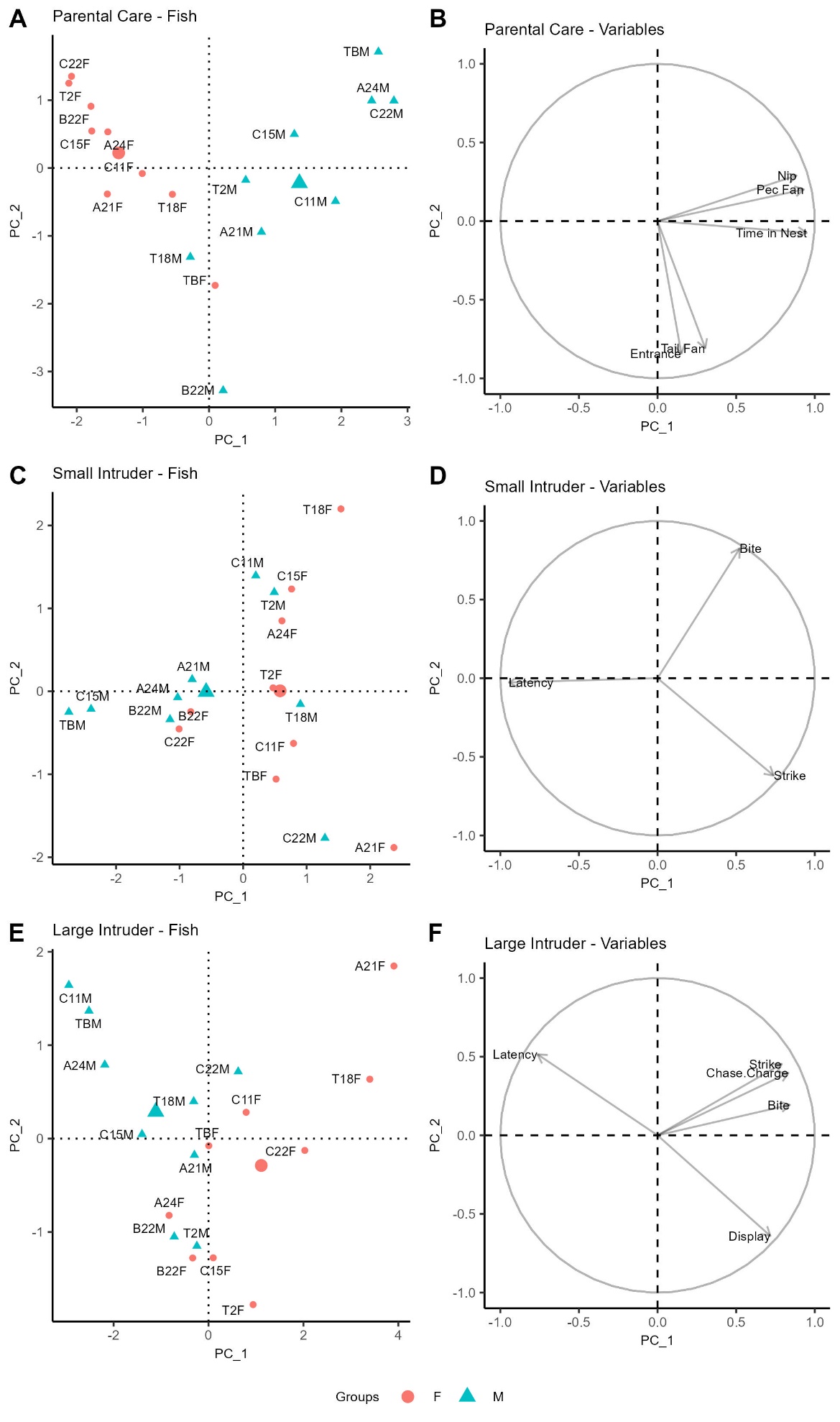

**Supplementary Figure 6. PCA for behavioral composition of parental care, small intruder aggression, and large intruder aggression.** For each figure, the left plot displays the loading of individuals based on PCs 1 and 2, and the right plot displays the variable loadings within each assay. Each PCA contains the mean of each behavior recorded in each assay collapsed across round and egg condition. In all plots, the larger blue triangle indicates the mean of males, and the larger red circle indicates the mean of females. A) Parental care PC1 vs PC2. Males separate from females based on any parental behavior. B) Small intruder aggression PC1 vs PC2. Males and females separate based almost exclusively based on latency, with males displaying higher latency. C) Large intruder aggression PC1 vs PC2. Males and females separated largely based on latency and displays, with males producing high latency and low displays, and females producing low latency and high displays.

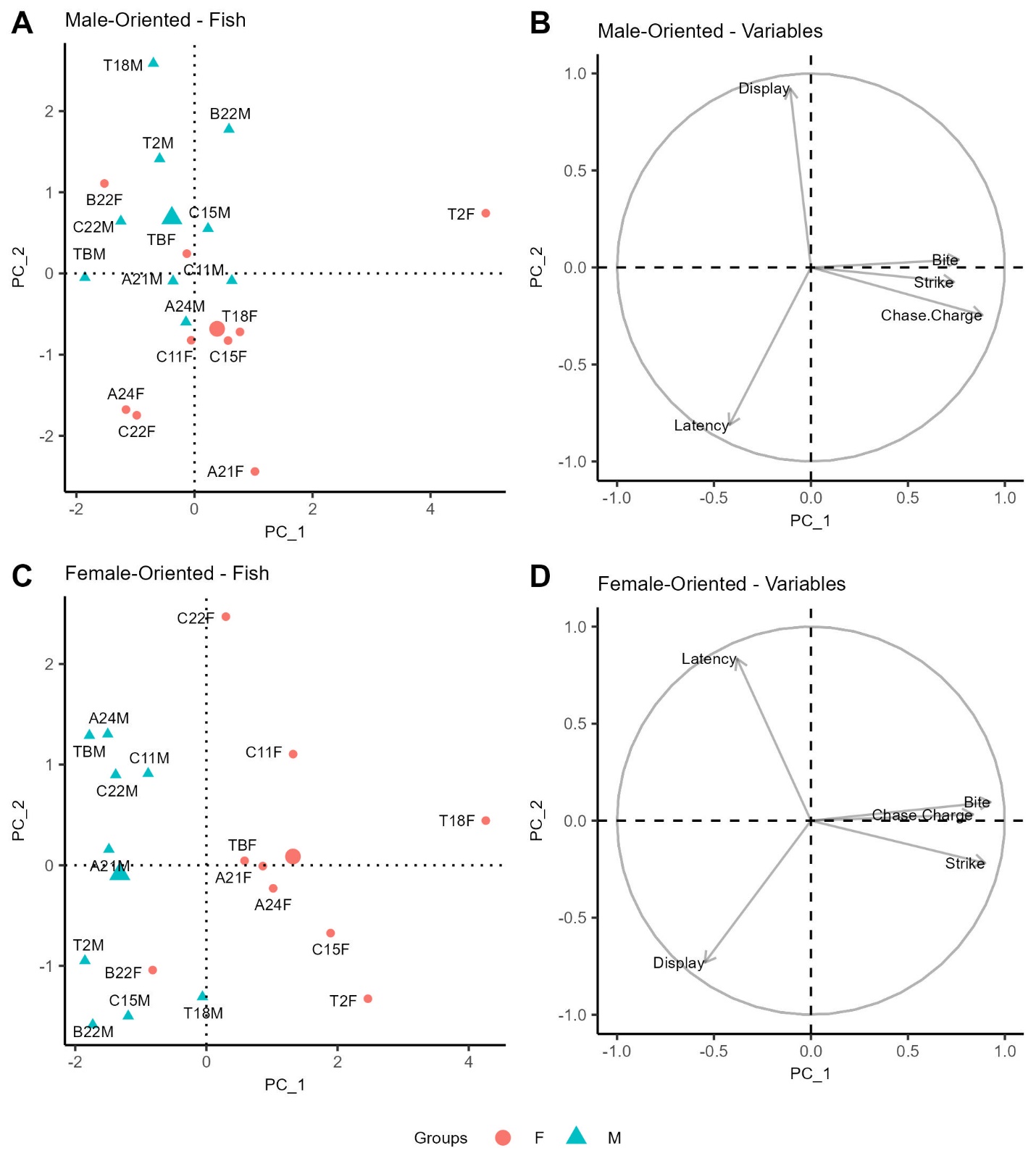

**Supplementary Figure 7. PCA for behavioral composition of male and female-oriented aggression.** For each figure, the left plot displays the loading of individuals based on PCs 1 and 2, and the right plot displays the variable loadings within each assay. Each PCA contains the mean of each behavior recorded in the assay collapsed across round and egg condition. In all plots, the larger blue triangle indicates the mean of males, and the larger red circle indicates the mean of females. A) Male-oriented aggression PC1 vs PC2. Males and females separate based on displays and latency, however unlike in other assays, males are characterized by high displays and low latency, with females displaying the opposite pattern. B) Female-oriented aggression PC1 vs PC2. Similar to in our statistical analysis, males and females separate based on direct aggression (bites, chases, strikes), and indirect aggression (displays) with males displaying high indirect aggression and low direct aggression, and females displaying the opposite pattern of aggression.

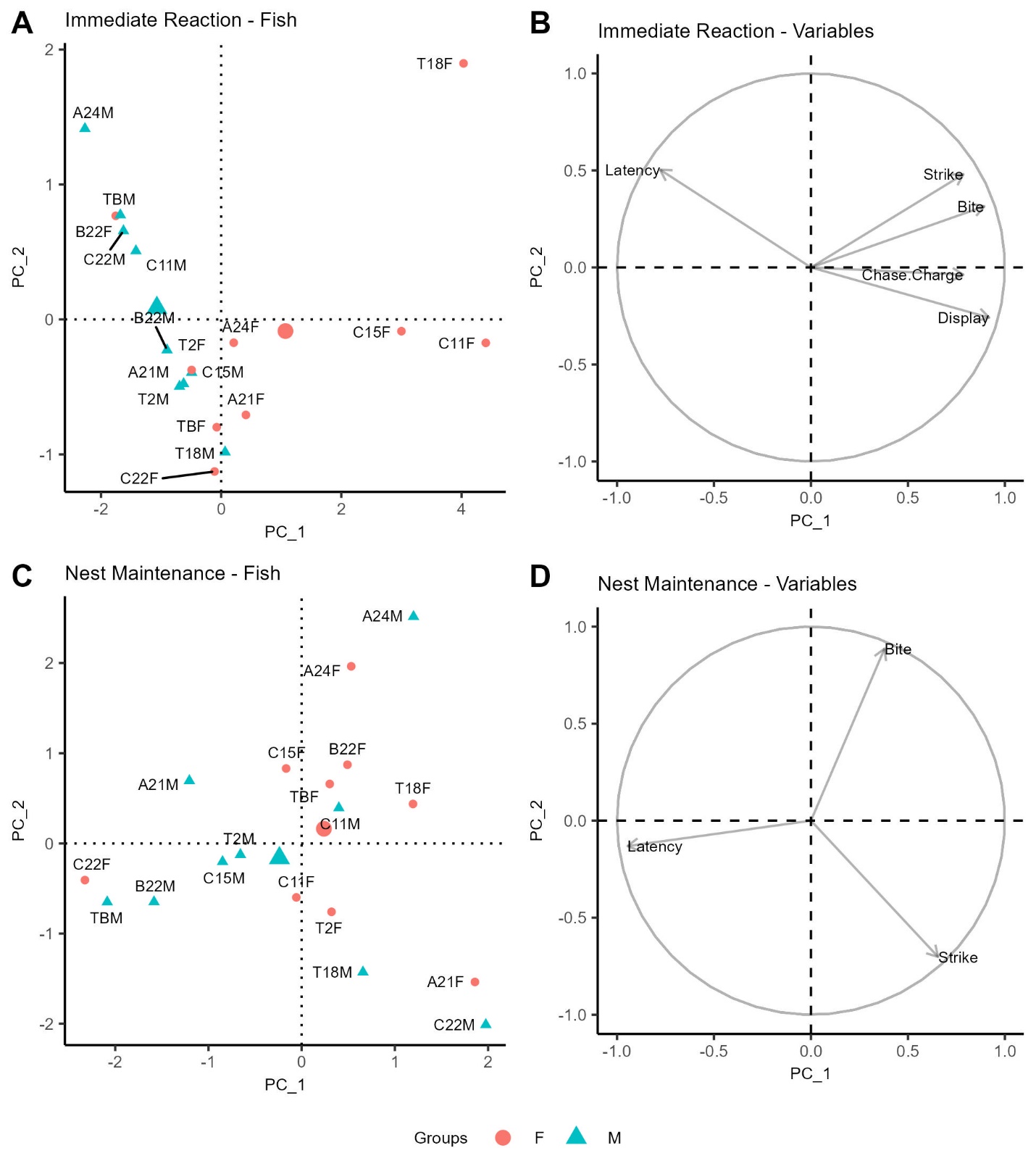

**Supplementary Figure 8. PCA for behavioral composition of immediate reaction to a threat and nest maintenance.** For each figure, the left plot displays the loading of individuals based on PCs 1 and 2, and the right plot displays the variable loadings within each assay. Each PCA contains the mean of each behavior recorded in an assay collapsed across round and egg condition. In all plots, the larger blue triangle indicates the mean of males, and the larger red circle indicates the mean of females. A) Immediate reaction to a threat. Males and females separate based on latency and aggression, with males being less aggressive and producing higher latency than females. B) Nest maintenance. Males and females do not separate particularly well, however males produce slightly higher latency, while females produce slightly more bites and strikes.

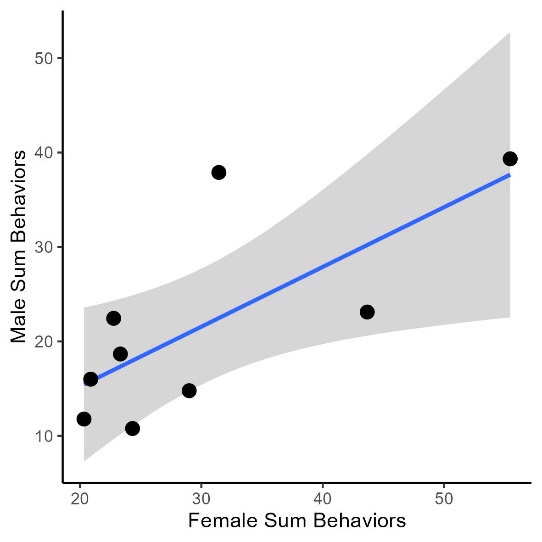

**Supplementary Figure 9**. **Male and female levels of aggression toward a female intruder are correlated.** The mean of the sum of aggressive behaviors recorded for females against the sum of aggressive behaviors recorded for males in female-oriented aggression collapsed across egg condition and round. Females’ behavior significantly predicts males’ behavior. The blue line represents simple linear regression lines. Ribbons show 95% confidence interval.
